# A brain network supporting social influences in human decision-making

**DOI:** 10.1101/551614

**Authors:** Lei Zhang, Jan P. Gläscher

## Abstract

Humans learn from their own trial-and-error experience and from observing others. However, it remains unanswered how brain circuits compute expected values when direct learning and social learning coexist in an uncertain environment. Using a multi-player reward learning paradigm with 185 participants (39 being scanned) in real-time, we observed that individuals succumbed to the group when confronted with dissenting information, but increased their confidence when observing confirming information. Leveraging computational modeling and fMRI we tracked direct valuation through experience and vicarious valuation through observation, and their dissociable, but interacting neural representations in the ventromedial prefrontal cortex and the anterior cingulate cortex, respectively. Their functional coupling with the right temporoparietal junction representing instantaneous social information instantiated a hitherto uncharacterized social prediction error, rather than a reward prediction error, in the putamen. These findings suggest that an integrated network involving the brain’s reward hub and social hub supports social influence in human decision-making.

## INTRODUCTION

Human decision-making is affected by direct experiential learning and social observational learning. This concerns both big and small decisions alike: In addition to our own experience and expectation, we care about what our family and friends think of which major we choose in college, and we also monitor other peoples’ choices at the lunch counter in order to obtain some guidance for our own menu selection—a phenomenon known as social influence. Classic behavioral studies have established a systematic experimental paradigm of assessing social influence^1^, and neuroimaging studies have recently attempted to unravel their neurobiological underpinnings^2, 3^. However, social influence and subsequent social learning^4^ has rarely been investigated in conjunction with direct learning.

Direct learning has been characterized in detail with reinforcement learning^5^ (RL) that describe action selection as a function of valuation, which is updated through a reward prediction error (RPE) as a teaching signal^6^. While social learning has been modeled by similar mechanism insofar as it simulates vicarious valuation processes of observed others^7, 8^, most studies only involved one single observed individual, and paradigms and corresponding computational models have not adequately addressed the aggregation of multiple social partners.

Albeit the computational distinction between direct learning (with experiential reward) and social learning (with vicarious reward), neuroimaging studies remain equivocal about the involved brain networks: Are neural circuits recruited for social learning similar to those for direct learning? In direct learning, a plethora of human functional magnetic resonance imaging (fMRI) studies have implicated a network involving the ventromedial prefrontal cortex (vmPFC) that represents individuals’ own valuation^9^, and the nucleus accumbens (NAcc) that encodes the RPE^10^. These findings mirror neurophysiological recordings in non-human primates showing the involvement of the orbitofrontal cortex and the striatum in direct reward experience^11, 12^. Turning to social learning, evidence from human neuroimaging studies have suggested similar neuronal patterns of experience-derived and observation-derived valuation, showing that the vmPFC processes values irrespective of being delivered to oneself or others^7, 13, 14^. However, recent studies in both human^15, 16^ and non-human primates^17, 18^ have suggested cortical contributions from the anterior cingulate cortex (ACC) that specifically tracks rewards allocated to others.

Intriguingly, although these findings suggest that direct learning and social learning are in part instantiated in dissociable brain networks, only very few studies have investigated how these brain networks interact when direct learning and social learning coexist in an uncertain environment^19^ and none of them involved groups larger than two individuals.

Here, we investigate the interaction of direct learning and social learning at behavioral, computational, and neural levels. We hypothesize that individuals’ direct valuation is computed via RL and has its neural underpinnings in the interplay between the vmPFC and the NAcc, whereas individuals’ vicarious valuation is updated by observing their social partners’ performance and is encoded in the ACC. In addition, we hypothesize that instantaneous socially based information has its basis in the right temporoparietal junction (rTPJ) that encodes others’ intentions necessary for choices in social contexts^15, 20, 21^. To test these hypotheses, we designed a multi-stage group decision-making task in which instantaneous social influence was directly measured as a response to the revelation of the group’s decision in real-time. By further providing reward outcomes to all individuals we enabled participants to learn directly from their own experience and vicariously from observing others. Our computational model updates direct and vicarious learning separately, but they jointly predict individuals’ decisions. Using model-based fMRI analyses we investigate crucial decision variables derived from the model, and through connectivity analyses, we demonstrate how different brain regions involved in direct and social learning interact and integrate social information into valuation and action selection. In addition, confidence was measured before and after receiving social information, as confidence may modulate individuals’ choices in social contexts^22, 23^.

Our data and model suggest that instantaneous social information alters both choice and confidence. After receiving outcome, experience-derived values and observation-derived values entail comparable contributions to inform future decisions but are distinctively encoded in the vmPFC and the ACC. We further identify an interaction of two brain networks that separately process reward information and social information, and their functional coupling substantiates a reward prediction error and a social prediction error as teaching signals for direct learning and social learning.

## RESULTS

Participants (N = 185) in groups of five performed the social influence task, of which, 39 were scanned with the MRI scanner. The task design utilized a multi-phase paradigm, enabling us to tease apart every crucial behavior under social influence (Fig. 1a). Participants began each trial with their initial choice (Choice 1) between two abstract fractals with complementary reward probabilities (70% and 30%), followed by their first post-decision bet^24^ (Bet 1, an incentivized confidence rating from 1 to 3). After sequentially uncovering the other players’ first decisions in the sequential order of participants’ subjective preference (i.e., participants decided on whose choice to see in the first and the second place, followed by the remaining two choices), participants had the opportunity to adjust their choice (Choice 2) and bet (Bet 2). The final choice and bet were then multiplied to determine the outcome on that trial (e.g., 3 × 20 = 60 cents). Participants’ actual choices were communicated in real time to every other participant via intranet connections, thus maintaining a high ecological validity. Importantly, the core of this paradigm was a probabilistic reversal learning task^25^ (PRL). This PRL implementation required participants to learn and continuously re-learn action-outcome associations, thus creating enough uncertainty such that group decisions were likely to be taken into account for behavioral adjustments in second decisions (before outcome delivery; referred to as instantaneous social influence), and for making future decisions on the next trial by observing others’ performance (after outcome delivery; i.e., social learning) together with participants’ own valuation process (i.e., direct learning). These dynamically evolving group decisions also allowed us to parametrically test the effect of group consensus, which moved beyond using only one social partner or an averaged group opinion^2, 23, 26^. Although participants were able to gain full action-outcome association at the single-trial level, across trials, participants may acquire additional valuation information by observing others, given the multiple reversal nature of the PRL paradigm. Additionally, participants were aware that there was neither cooperation nor competition (Methods).

**Fig. 1.**
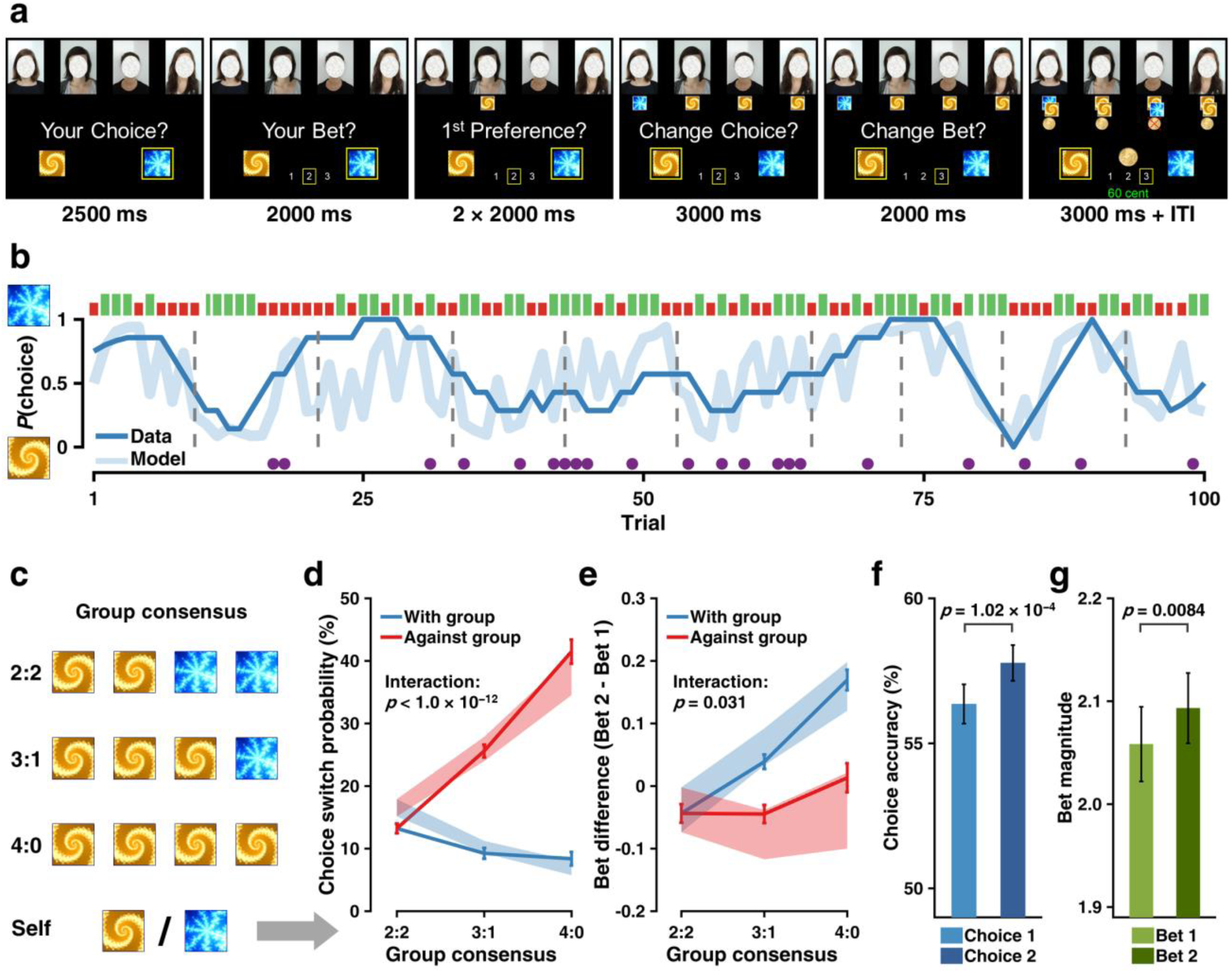
Experimental task and behavioral results. **(a)** Task design. Participants (N = 185) made an initial choice and an initial bet (Choice 1, Bet 1), and after observing the other four co-players’ initial choices, they were asked to adjust their choice and bet (Choice 2, Bet 2), followed by the outcome. **(b)** Example task dynamic. Trial-by-trial behavior for an example participant. Blue curves, seven-trial running averages of choices (dark) and predicted choice probabilities from the winning model M6b (light). Green (long) and red (short) bars, rewarded and unrewarded trials; purple circles, switches on Choice 2; dashed vertical lines, reversals that took place every 8–12 trials. **(c)** Illustration of group consensus (view from each participant). **(d)** Social influence on choice adjustments. Choice switch probability as a function of group consensus, illustrated in (c), and direction (with vs. against) of the majority of the group. Results indicated a main effect of direction (*F*_1,228_ = 299.63, *p* < 1.0 × 10^−15^), a main effect of consensus (*F*_2,574_ = 131.49, *p* < 1.0 × 10^−15^), and an interaction effect (*F*_1,574_ = 55.82, *p* < 1.0 × 10^−12^). Solid lines indicate actual data (mean ± within-subject standard error of the mean, SEM). Shaded error bars represent the 95% highest density interval (HDI) of mean effects computed from the winning model M6b’s posterior predictive distribution. **(e)** Social influence on bet adjustments. Bet difference as a function of group consensus and direction of the majority of the group. Results indicated a main effect of direction (*F*_1,734_ = 50.95, *p* < 1.0 × 10^−11^), a main effect of consensus (*F*_2,734_ = 16.74, *p* < 1.0 × 10^−7^), and an interaction effect (*F*_1,734_ = 4.67, *p* =0.031). Format is as in Fig. 1d. **(f–g)** Enhanced performance after adjustment. (f) Accuracy of Choice 2 was higher than that of Choice 1 (*t*185 = 3.971, *p* = 1.02 × 10^−4^). (g) Magnitude of Bet 2 was larger than that of Bet 1 (*t*185 = 2.665, *p* = 0.0084).

### Instantaneous Social Influence Alters Both Action and Confidence in decision-making

Human participants’ choices tracked option values over probabilistic reversals (Fig. 1b). Interestingly, participants indeed changed their choice and bet after observing group decisions, but in the opposite direction. Both the choice adjustment and the bet adjustment were modulated by a significant interaction between the relative direction of the group (with vs. against) and the group consensus (2:2, 3:1, 4:0, view of each participant, Fig. 1c). In particular, participants showed an increasing trend to switch their choice toward the group when faced with more dissenting social information, whereas, they were more likely to persist when observing agreement with the group (direction × consensus: *F*_1,574_ = 55.82, *p* < 1.0 × 10^−12^; Fig. 1d). Conversely, participants tended to increase their bets as a function of the group consensus when observing confirming opinions, but sustained their bets when being contradicted by the group (direction × consensus: *F*_1,734_ = 4.67, *p* = 0.031; Fig. 1e). Bet difference was also analyzed conditioned on participants’ switching behavior on Choice 2, and results were in coherent with the main findings (Supplementary Fig. 2a).

We further verified the benefit of considering instantaneous social information for behavior adjustments. Participants’ choice accuracy of the second decision was significantly higher than that of the first one (*t*_185_ = 3.971, *p* = 1.02 × 10^−4^; Fig. 1f; Supplementary Fig. 2b), and participants’ second bet was significantly larger than their first one (*t*_185_ = 2.665, *p* = 0.0084; Fig. 1g, Supplementary Fig. 2c). These results suggested that, in the case of behavioral adjustments, despite that participants were often confronted with conflicting group decisions, considering social information in fact facilitated learning. Notably, these behavioral adjustments were not likely due to perceptual conflict, in which participants would have made switches in a random fashion, hence no learning enhancement. Strikingly, no such benefit of adjustment was observed in a non-social control experiment, where participants (N = 36; Supplementary Note 1) were performing this task with intelligent computer agents (Supplementary Fig. 1a–f). It is worth noting that although we did not intentionally manipulate the amount of dissenting social information (given the real-time property), it was nonetheless randomly distributed (*p*s > 0.05, Wald-Wolfowitz test). Moreover, neither the amount of dissenting social information nor participants’ choice switching behavior was related to the time of reversal or the lapse error indicated by our winning model (Methods; Supplementary Fig. 2d,e).

Taken together, our behavioral results demonstrated that instantaneous social information altered individuals’ choice and confidence, which accounted for facilitated learning after behavioral adjustment, and this benefit could not be explained by perceptual mismatch and may be specific only when interacting with human partners.

### Computational Mechanisms of Integrated Valuation Between Direct Learning and Social Learning

Using computational modeling, we aimed to formally quantify latent mechanisms that underlay the learning processes in our task on a trial-by-trial basis. Different from existing RL models on social influence^26^, our model accommodates multiple players and is able to simultaneously estimate all participants’ behaviors (both choices and both bets) under the hierarchical Bayesian analysis workflow^27^. Our efforts to construct the winning model (Fig. 2a) were guided by two design principles: (1) separating individual’s own valuation updated via direct learning from vicarious valuation updated via social learning; (2) distinguishing instantaneous social influence before outcomes were delivered from social learning in which action-outcome associations were observed from the others. These design principles tied closely with our multiple task phases, representing a computationally plausible information flow.

**Fig. 2.**
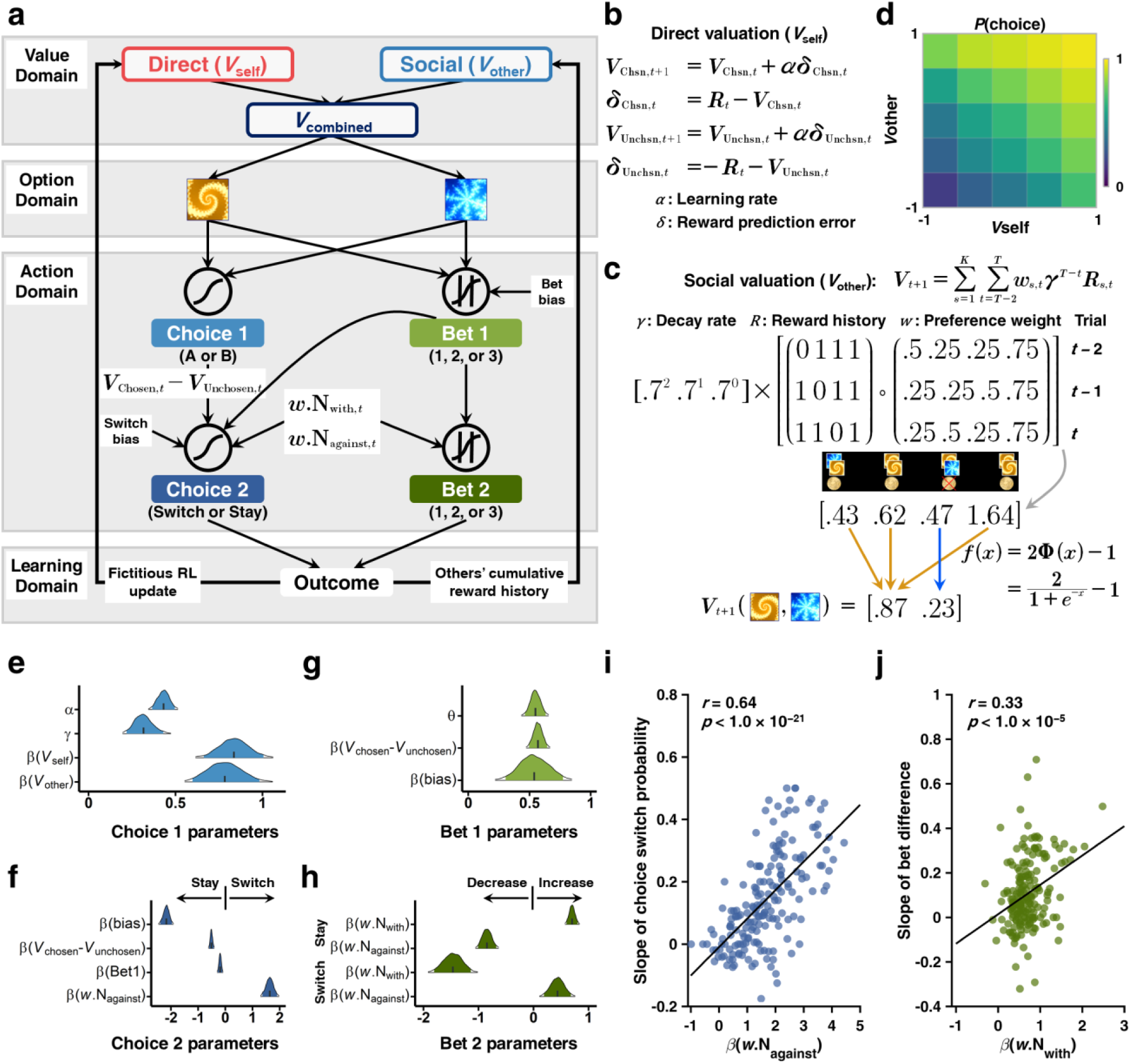
Computational model and its relation to behavior. **(a)** Schematic representation of the winning computational model (M6b). Participants’ initial behaviors (Choice 1, Bet 1) were accounted for by value signals updated from both direct learning and social learning; behavioral adjustments (Choice 2 and Bet 2) were ascribed to the valuation of initial behaviors (*V*_chosen,*t*_ – *V*_unchosen,*t*_) and preference-weighted instantaneous social information; value from direct learning (*V*_self_) was updated via a fictitious reinforcement learning model, while value from social learning (*V*_other_) was updated through tracking other co-players’ cumulative reward histories, weighted by preference and a decay rate. **(b)** Computation of *V*_self_. *V*_self_ was computed with a fictitious update reinforcement learning model, where values of both the chosen and the unchosen options were updated. **(c)** Computation of *V*_other_. *V*_other_ was computed as an exponentially decayed and preference-weighted other co-players’ cumulative reward histories in the last three trials (*t*-2 to *t*), normalized to lie between −1 and 1. **(d)** Contribution of *V*_self_ and *V*_other_ to action probability of Choice 1. Both *V*_self_ and *V*_other_ spanned within their range (−1 to 1), and they jointly contributed to *p*(Choice 1). **(e–h)** Model parameters of the winning model M6b. Posterior density for parameters related to Choice 1 (e), Choice 2 (f), Bet 1 (g), and Bet 2 (h). Short vertical bars indicate the posterior mean. Shaded areas depict 95% of the highest density interval (HDI). **(i–j)** Relationship between model parameters and behavioral results. (i) Relationship between dissenting social information (*w*.N_against_) and the susceptibility to social influence (i.e., slope of switch probability calculated from Fig. 1d; *r* = 0.64, *p* < 1.0 × 10^−21^). (j) Relationship between confirming social information (*w*.N_with_) and the extent of bet difference (i.e., slope of bet difference calculated from Fig. 1E; *r* = 0.33, *p* < 1.0 × 10^−5^).

On each trial, the option value of Choice 1 (A or B) was modeled as a linear combination between values from direct learning (*V*_self_) and values from social learning (*V*_other_):

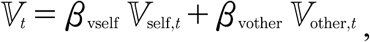

Where

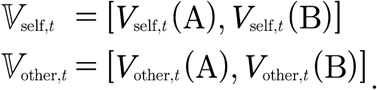

After participants discovered the other co-players’ first choices, their Choice 2 (switch or stay) was modeled as a function of two counteracting influences: (a) the preference-weighted group dissension (*w*.N_against_) representing the instantaneous social influence and (b) the difference between participants’ action values of Choice 1 (V_chosen,C1,*t*_ – V_unchosen,C1,*t*_) representing the distinctiveness of the current value estimates.

Lastly, when all outcomes were delivered, both *V*_self_ and *V*_other_ were updated. Notably, *V*_self_ was updated using the fictitious Rescorla-Wagner RL model^28^ (Fig. 2b), whereas *V*_other_ was updated through tracking an exponentially decayed and preference-weighted all four other co-players’ cumulative reward histories (i.e., their performance in the recent past; Fig. 2c). It is worth noting that our construction of *V*_other_ was in close accordance with previous evidence that suggested a discounted outcome history contributes to animals’ valuation processes^29^, and that the construction of *V*_other_ depicted social learning by simulating a vicarious valuation process by observing others^4, 16, 21, 30^. More importantly, the social learning here was weighted by social preference (*w_s,t_*) that reflected credibility assignment based on the social partners’ performance^15, 21^. Intriguingly, *V*_other_ did not contribute to the learning performance in the non-social control task despite similar behavioral adjustment patterns compared to the main study, suggesting the uniqueness of social learning in social contexts (Supplementary Fig. 1f). Together, all the above properties granted the social feature of *V*_other_ and demonstrated its distinct contribution in addition to *V*_self_.

We tested the winning model against alternative computational hypotheses under the hierarchical Bayesian framework^27^ (Table 1). We further verified our winning model using two rigorous validation approaches. First, we carried out a parameter recovery analysis to assure all parameters could be accurately and selectively identified (Supplementary Fig. 3 and Note 3).

**Table 1.** Candidate computational models and model comparison

| Class | Model | Description | # Par. | ΔLOOIC | Weight |
| --- | --- | --- | --- | --- | --- |
| Non-social models | M1a | Simple Rescorla-Wagner RL | 9 | 3614 | 0 |
|  | M1b | Fictitious RL | 9 | 2369 | 0 |
|  | M1c | Pearce-Hall | 11 | 8540 | 0 |
| Social models:<br>instantaneous<br>social influence | M2a | M1a + instantaneous social influence | 9 | 1721 | 0 |
|  | M2b | M1b + instantaneous social influence | 9 | 725 | 0 |
|  | M2c | M1c + instantaneous social influence | 11 | 2715 | 0 |
| Social models:<br>instantaneous<br>social influence<br>and social learning | M3 | M2b + SL (others' RL update) | 15 | 535 | .002 |
|  | M4 | M2b + SL (others' action preference) | 13 | 745 | 0 |
|  | M5 | M2b + SL (others' current reward) | 13 | 411 | 0 |
|  | M6a | M2b + SL (others' cumulative reward) | 14 | 164 | 0 |
|  | **M6b** | M2b + SL (others' cumulative reward) + bet1 | 15 | 0 | .998 |

Note: RL = reinforcement learning; SL = social learning; # Par. = number of free parameters at the individual level; ΔLOOIC = leave-one-out information criterion relative to the winning model (lower LOOIC value indicates better out-of-sample predictive accuracy); Weight = model weight calculated with Bayesian model averaging using Bayesian bootstrap (higher model weight value indicates higher probability of the candidate model to have generated the observed data). M6b (in bold) is the winning model.

Second, as model comparison provided relative model performance, we noted the importance to perform posterior predictive checks (PPC), and we found that the posterior prediction well captured key behavioral patterns (Fig. 1d,e, Supplementary Fig. 2a).

Parameter estimation results (Fig. 2e–h) suggested that the extent to which participants learned from themselves and from the others was on average comparable (*β*(*V*_self_) = 0.84, 95% HDI: [0.67, 1.01]; *β*(*V*_other_) = 0.78, 95% HDI: [0.59, 0.97]), suggesting value signals computed from direct learning and social learning were jointly employed to guide future decisions. Furthermore, parameters related to instantaneous social information were well-capable of predicting individual differences of participants’ behavioral adjustment: If the model-derived signal was in high accordance with the corresponding pattern of behavioral adjustment, we ought to anticipate a strong association between them. Indeed, we observed a positive correlation between *β*(*w*.N_against_) and slopes of choice switch probabilities in the against condition (*r* = 0.64, *p* < 1.0 × 10^−21^; Fig. 2i; slopes computed from Fig. 1d). Similarly, we observed a positive correlation between *β*(*w*.N_with_) and slopes derived from bet differences in the “with” condition (*r* = 0.33, *p* < 1.0 × 10^−5^; Fig. 2j; slopes computed from Fig. 1e). Taken together, our computational modeling analyses suggested that participants learned both from their direct valuation process and from vicarious valuation experience, and values from direct learning and social learning jointly contributed to the decision process. Moreover, participants’ behavioral adjustments were predicted by the counteracting effects between their initial valuation and the instantaneous social information. Next, once we had uncovered those latent variables of the decision processes underlying the social influence task, we were able to test how they were computed and implemented at the neural level using model-based fMRI^31^.

### Neural Substrates of Dissociable Value Signals from Direct Learning and Social Learning

The first part of our model-based fMRI analyses focused on how distinctive decision variables (Fig. 3a) were represented in the brain (GLM 1). We aimed to test the hypothesis that distinct and dissociable brain regions were recruited to implement direct learning and social learning signals (i.e., component value^22^). We observed that the vmPFC (see Table 2 for all MNI coordinates and multiple comparisons correction methods) activity was positively scaled with *V*_self_, and the ACC activity was positively scaled with *V*_other_ (Fig. 3b). To test whether the two value signals were distinctively associated with vmPFC and ACC, we employed a double-dissociation approach, and we found that *V*_self_ was exclusively encoded in the vmPFC (*β* = 0.1458, *p* < 1.0 × 10^−5^; Fig. 3e, red) but not in the ACC (*β* = 0.0128, *p* = 0.4394; Fig. 3d, red), whereas *V*_other_ was exclusively represented in the ACC (*β* = 0.1560, *p* < 1.0 × 10^−5^; Fig. 3d, blue) but not in the vmPFC (*β* = 0.0011, *p* = 0.9478; Fig. 3e, blue). Computationally, these two sources of value signals needed to be integrated to make decisions (i.e., integrated value^22^). We reasoned that if a region is implementing the integrated value, it must have functional connectivity with regions tracking each of the value signals (i.e., vmPFC, ACC). Using a physio-physiological interaction analysis, we found that the medial prefrontal cortex covaried with both the vmPFC and the ACC (Supplementary Fig. 6a).

**Fig. 3.**
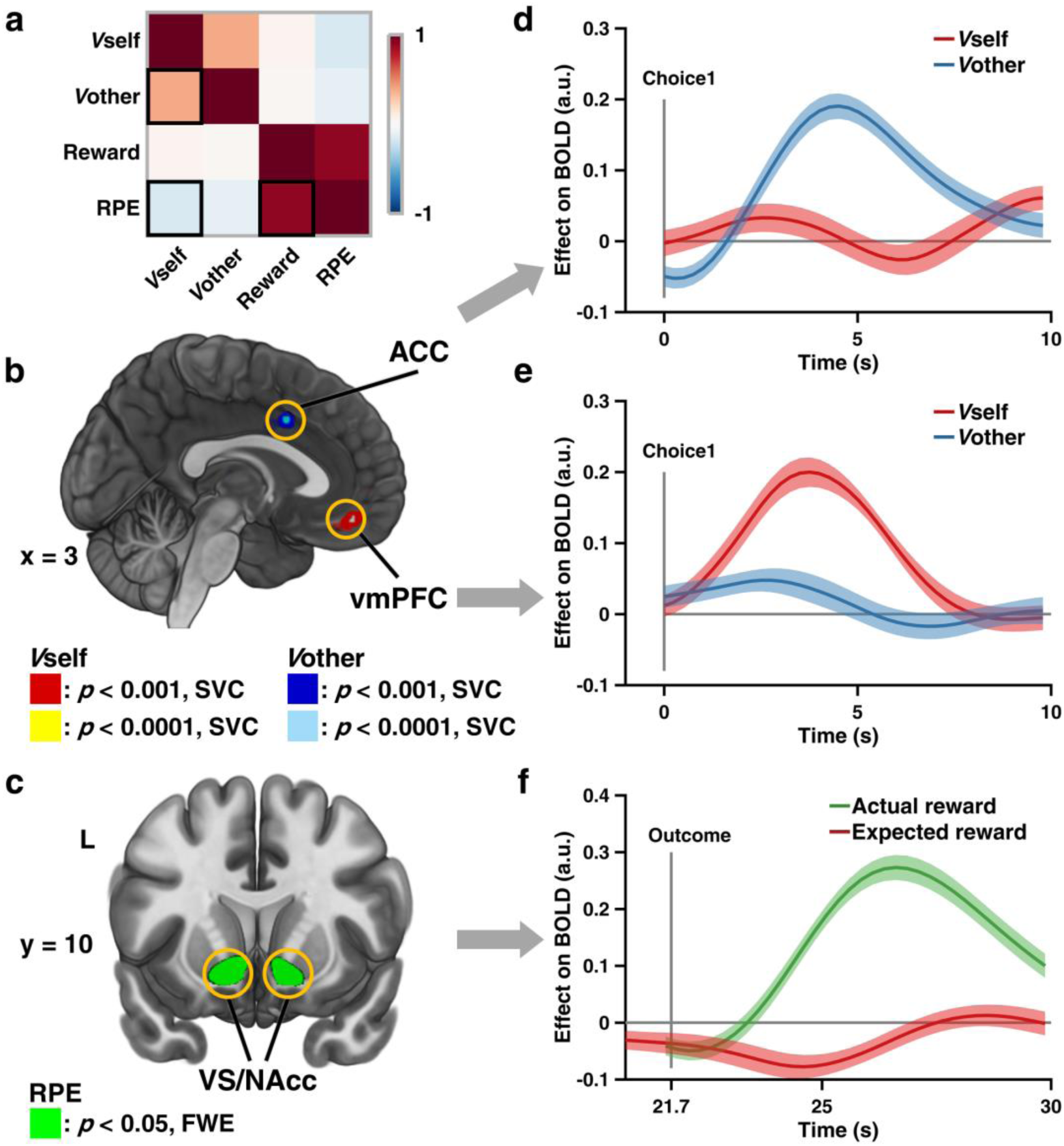
Neural substrates of dissociable value signals and reward prediction error. **(a)** Correlation matrix of value-related decision variables derived from M6b. **(b)** Neural representation of value signals. *V*_self_ and *V*_other_ were encoded in the vmPFC (red/yellow) and the ACC (blue/light blue), respectively. Display thresholded at *p* < 0.001 and *p* < 0.0001, small volume corrected (SVC); sagittal slice at x = 3. Actual results were TFCE SVC-corrected at *p* < 0.05. **(c)** Neural representation of reward prediction error (RPE). RPE was encoded in the VS/NAcc. Display thresholded at *p* < 0.05, family-wise error (FWE) corrected; coronal slice at y = 10. Actual results were TFCE whole-brain FWE corrected at *p* < 0.05. **(d–e)** ROI time series analyses of vmPFC and ACC, demonstrating a double dissociation of the neural signatures of value signals. (d) BOLD signal of ACC was only positively correlated with *V*_other_ (*β* = 0.1560, *p* < 1.0 × 10^−5^, permutation test; blue line), but not with *V*_self_ (*β* = 0.0011, *p* = 0.9478, permutation test; red line), whereas (e) BOLD signal of vmPFC was only positively correlated with *V*_self_ (*β* = 0.1458, *p* < 1.0 × 10^−5^, permutation test; red line), but not with *V*_other_ (*β* = 0.0128, *p* =0.4394, permutation test; blue line). Lines and shaded areas show mean ± SEM of *β* weights across participants. **(f)** ROI the time series analyses of VS/NAcc, showing its sensitivity to both components of RPE (i.e., actual reward and expected reward). BOLD signal of VS/NAcc was positively correlated with actual reward (*β* = 0.2298, *p* < 1.0 × 10^−5^, permutation test; green line), and negatively correlated with expected reward (*β* = −0.0327, *p* = 0.021, permutation test; red line). Format is as in Fig. 3d.

**Table 2.**
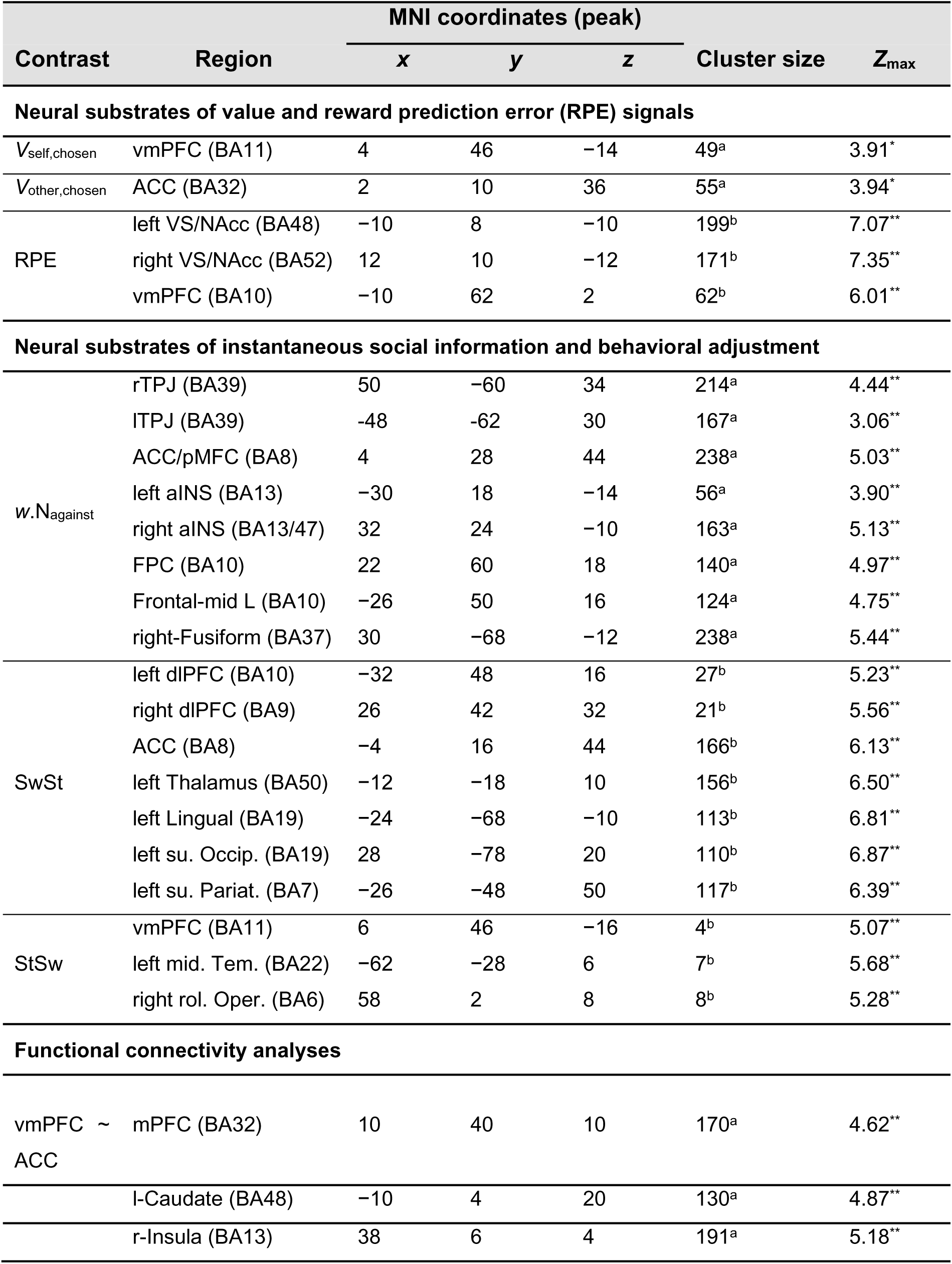
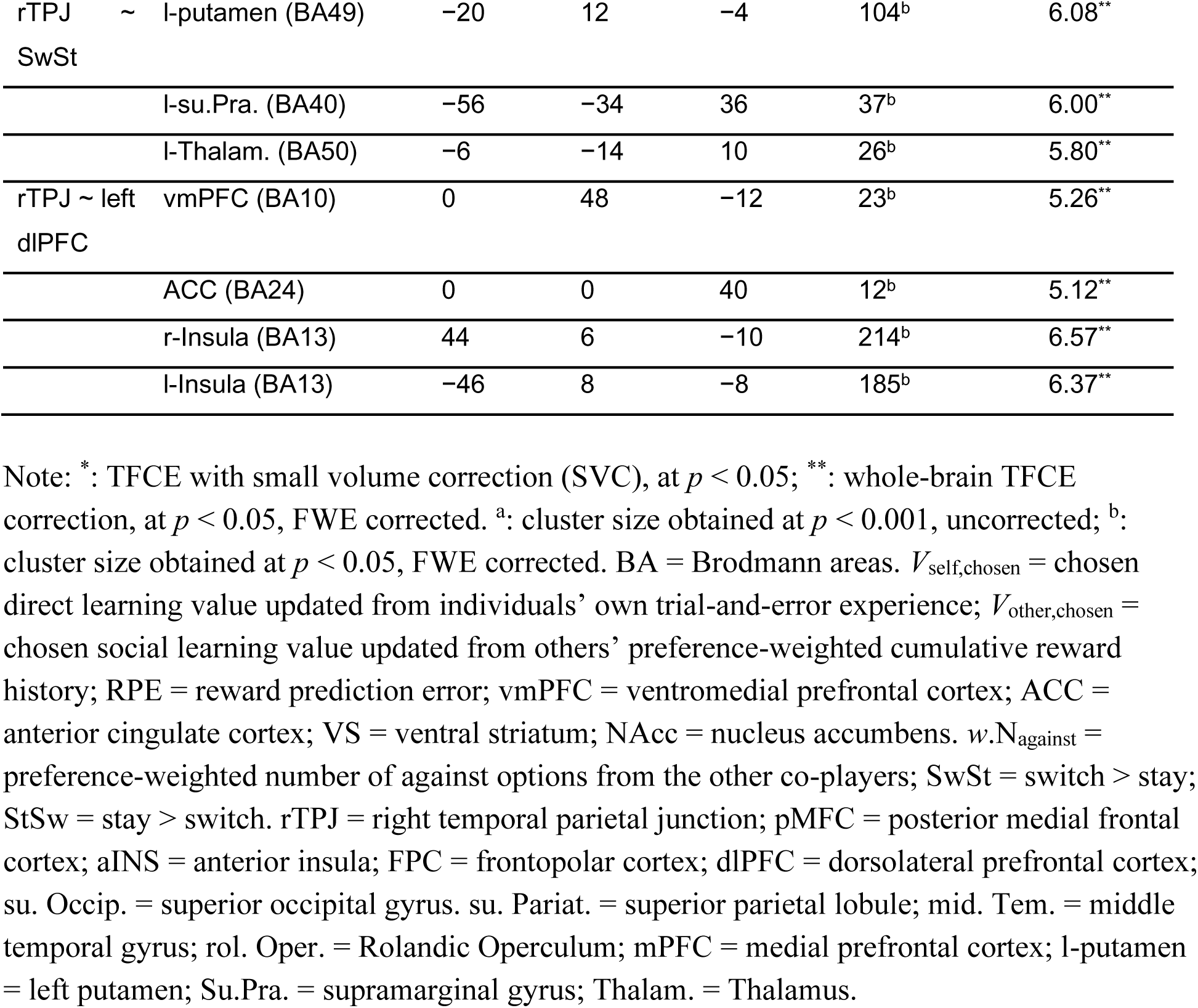
Neural substrates of decision variables.

Besides the value signals, the RPE signal was firmly associated with activities in the bilateral NAcc (Fig. 3c). Furthermore, a closer look at the two theoretical sub-components of RPE was necessary to assess its neural substrates^15, 32^. Specifically, according to the specification of RPE (Fig. 2b), to qualify as a region encoding the RPE signal, activities in the NAcc ought to covary positively with the actual outcome (i.e., reward) and negatively with the expectation (i.e., value). This property thus provides a common framework to test the neural correlates of any error-like signal. Under this framework, we indeed found that activities in the NAcc showed a positive effect of the reward (*β* = 0.2298, *p* < 1.0 × 10^−5^), and a negative effect of the value (*β* =−0.0327, *p* = 0.021; Fig. 3f), justifying that NAcc was encoding the RPE signal instead of the outcome valence. Variables related to participants’ bet did not yield significant clusters.

### Neural Correlates of Dissenting Social Information and Behavioral Adjustment

We next turned to disentangle the neural substrates of the instantaneous social influence (GLM 1) and the subsequent behavioral adjustment (GLM 2). Since we have validated enhanced learning after considering instantaneous social information (Fig. 1f–g), we reasoned that participants might process other co-players’ intentions relative to their own first decision to make subsequent adjustments, and this might be related to the mentalizing network. Based on this reasoning, we assessed the parametric modulation of preference-weighted dissenting social information (*w*.N_against_), and found that activities in the TPJ, among other regions (Table 2), were positively correlated with the dissenting social information (Supplementary Fig. 4). Furthermore, the resulting choice adjustment (i.e., switch > stay) covaried with activity in bilateral dorsolateral prefrontal cortex (Supplementary Fig. 5a,d), commonly associated with executive control and behavioral flexibility^25^. By contrast, the vmPFC was more active during stay trials (i.e., stay > switch), reminiscent of its representation of one’s own valuation (Supplementary Fig. 5d,f). Hence, these findings were not likely due to learning of the task structure, but rather, were genuinely attributed to dissenting social information and choice adjustment, respectively.

### A Network between Brain’s Reward Circuits and Social Circuits

Above we demonstrated how key decision variables related to value and reward processing and social information processing were implemented at distinct nodes at the neural level. In the next step, we sought to establish how these network nodes were functionally connected to bring about socially-induced behavioral change and to uncover additional latent computational signals that would otherwise be undetectable by conventional general linear models.

Using a psycho-physiological interaction (PPI), we investigated how behavioral change at Choice 2 was associated with the functional coupling between rTPJ that processed instantaneous social information and other brain regions. This analysis identified enhanced connectivity between left putamen (Fig. 4a–c) and rTPJ as a function of choice adjustment. Closer investigations into the computational role of lPut revealed that it did not correlate with both sub-components of the RPE (Supplementary Fig. 6c). Instead, as the choice adjustment resulted from processing social information, we reasoned that lPut might encode a social prediction error (SPE) at the time of observing social information, delineating the difference between the actual consensus and the expected consensus of the group. Specifically, the expected consensus was approximated by the difference in participants’ vicarious valuation (*V*_other,chosen,*t*_ – *V*_other,unchosen,*t*_), on the basis that knowing how the others value specific options helps individuals model the others’ future behaviors^30, 33^ (e.g., when *V*_other,chosen,*t*_ – *V*_other_,unchosen,*t* was large, participants were relatively sure about option values learned from the others, therefore anticipating more coherent group choices). Following this reasoning, we conducted a similar time series analysis as we did for the RPE, and we found that activity in the lPut was indeed positively correlated with the actual consensus (*β* = 0.0363, *p* = 0.0438) and negatively correlated with the expected consensus (*β* = −0.0409, *p* = 0.0123; Fig. 4d). This pattern suggested that lPut was effectively encoding a hitherto uncharacterized social prediction error rather than a reward prediction error (Supplementary Fig. 6b). Taken together, these analyses demonstrated that the functional coupling between neural representations of social information and of SPE was enhanced, when this social information was leading to a behavioral change.

**Fig. 4.**
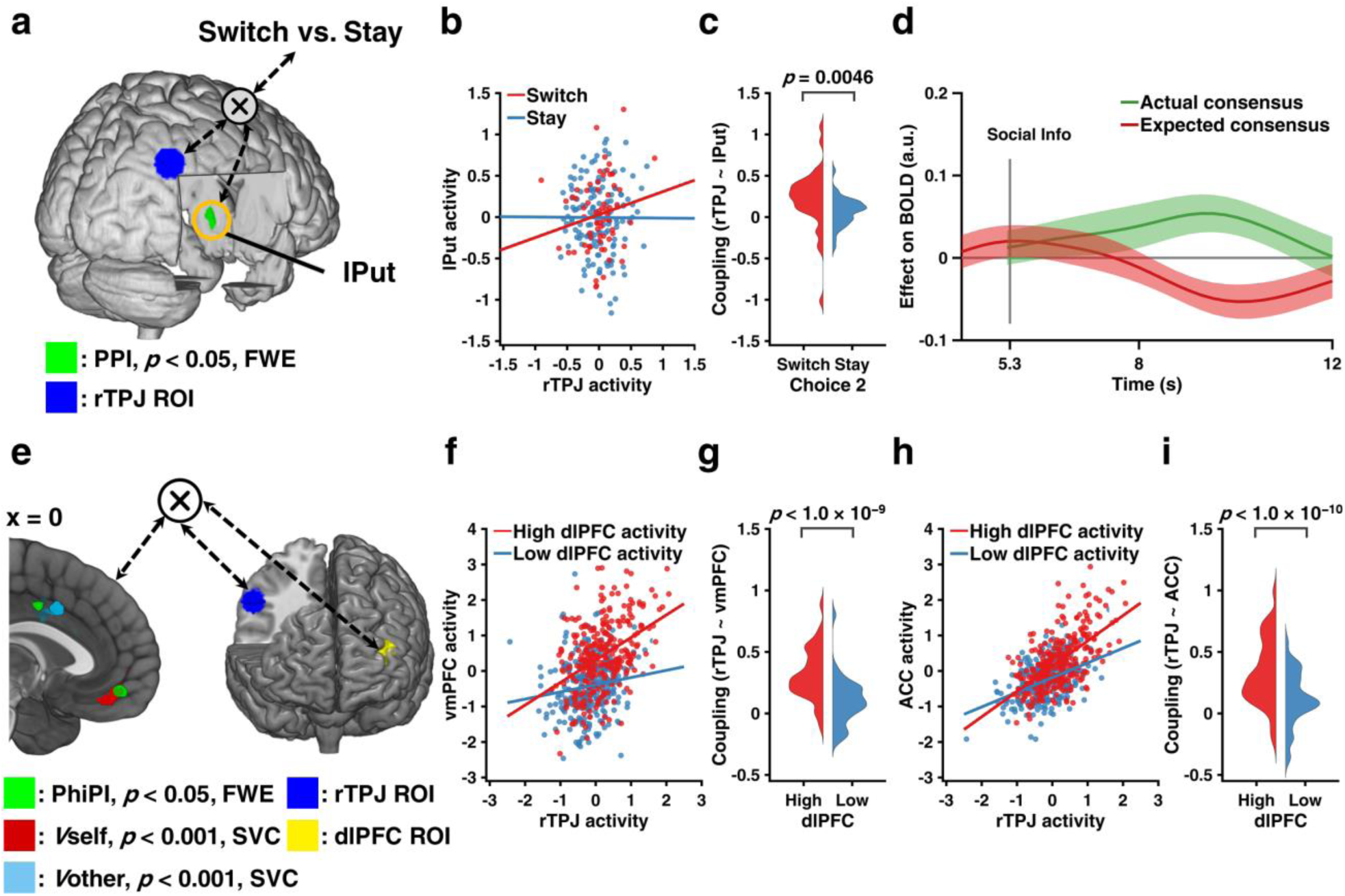
Functional connectivity between reward-related regions and social-related regions. **(a)** Increased functional connectivity between the left putamen (green) and the seed region rTPJ (blue) as a function of choice adjustment (switch vs. stay). Display thresholded at *p* < 0.05, FWE-corrected. Actual results were TFCE whole-brain FWE corrected at *p* < 0.05. **(b)** Correlation of activity in seed and target regions for both switch and stay trials in an example subject. **(c)** Kernel density estimation of coupling strength across all participants for switch and stay trials. **(d)** ROI time series analyses of the left putamen (lPut), exhibiting a social prediction error signal: BOLD signal of lPut was positively correlated with the actual consensus (*β* = 0.0363, *p* = 0.0438, permutation test; green line), and negatively correlated with the expected consensus (*β* = −0.0409, *p* = 0.0123, permutation test; red line). Format is as in Fig. 3d. **(e)** Physio-physiological interaction between social-related regions and reward-related regions. The rTPJ seed (blue) and the left dlPFC seed (yellow) elicited connectivity activations (target regions) in the vmPFC and the pMFC (both in green), which partially overlapped with neural correlates of value signals in vmPFC and ACC, as in Fig. 3b. Display thresholded at *p* < 0.05, FWE-corrected; sagittal slice at x = 0. Actual results were TFCE whole-brain FWE corrected at *p* < 0.05. **(f–i)** Correlation plots of seed and target regions for both high and low dlPFC activities in an example subject (f, h) and kernel density estimation of seed-target coupling strengths across all participants for high and low dlPFC activities (g, i).

In the last step, using a physio-physiological interaction (PhiPI) we investigated how neural substrates of switching at Choice 2 in the left dlPFC were accompanied by the functional coupling of rTPJ and other brain regions. This analysis revealed that rTPJ covaried with both vmPFC, scaled by the activation level of dlPFC (Fig. 4e–i). Strikingly, these target regions overlapped with regions that represented two value signals in vmPFC and ACC that we reported earlier (c.f., Fig. 3b). Collectively, our functional connectivity analyses suggested the interplay of brain regions representing social information and the propensity for behavioral change led to the neural activities of values signals in the vmPFC and ACC, which are updated via both direct learning and social learning.

## DISCUSSION

Social influence is a powerful modulator of individual choices, yet how social influence and subsequent social learning interact with direct learning in a probabilistic environment is poorly understood. Here, we bridge this gap with a multi-player social decision-making paradigm in real-time that allowed us to dissociate between experience-driven valuation and observation-driven valuation. In a comprehensive neurocomputational approach, we are not only able to identify a network of brain regions that represents and integrates social information in learning, but also characterize the computational role of each node in this network in detail (Fig. 5), suggesting the following process model: Individuals’ own decision is guided by a combination of value signals from direct learning (*V*_self_) represented in the vmPFC (Fig. 3b,e) and from social learning (*V*_other_) represented in a section of the ACC (Fig. 3b,d). The instantaneous social information reflected by decisions from others are encoded with respect to one’s own choice in the rTPJ (Supplementary Fig. 4), an area linked, but not limited to representations of social information and social agents in a variety of tasks^20, 34^. In fact, rTPJ is also related to Theory of Mind^35^ and other integrative computations such as multisensory integration^36^ and attentional processing^37^. Moreover, dissenting social information gives rise to a hitherto uncharacterized social prediction error (difference between actual and expected consensus of the group) represented in the putamen (Fig. 4d), unlike the more medial NAcc, which exhibits the neural signature of a classic reward prediction error^10^ (Fig. 3c,f). Notably, the interplay of putamen and rTPJ modulates behavioral change toward the group decision (Fig. 4a–c) in combination with its neural representation of choice switching in the dlPFC (Fig. 4e–i). These connected neural activations functionally couple with the valuation of direct learning in the vmPFC (*V*_self_) and social learning in the ACC (*V*_other_), thus closing the loop of decision-related computations in social contexts.

**Fig. 5.**
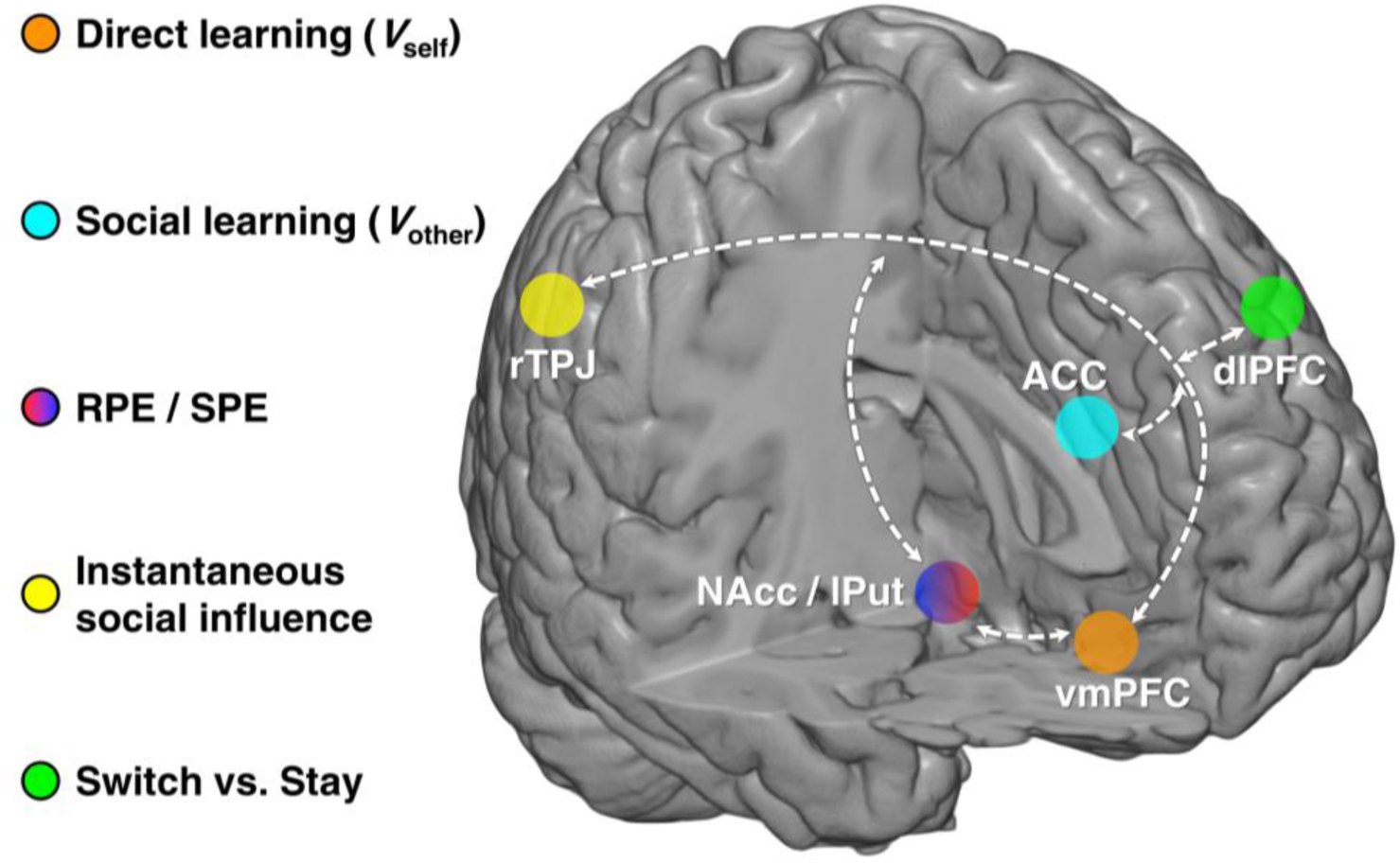
Schematic illustration of the brain network supporting social influence in decision-making as uncovered in this study **(for details see main text)**.

Our result that direct valuation is encoded in the vmPFC are firmly in line with previous evidence from learning and decision-making in non-social contexts^9^, and demonstrated the role of vmPFC in experiential learning into a social context. In addition to individuals’ own value update, we further show that the ACC encodes value signals updated from social learning, which is aligned with previous studies that have implicated the role of ACC in tracking the volatility of social information^15^ and vicarious experience^38^. In particular, given that social learning in the current study is represented by the preference-weighted cumulative reward histories of the others, the dynamics of how well the others were performing in the recent past somewhat reflects their volatility in the same learning environment^15^. Moreover, this distinct neural coding of direct values and vicarious values in the current study fundamentally differs from previous studies on social decision-making. While previous studies have found evidence for a role of vmPFC and ACC in encoding self-oriented and other-oriented information^39^, those signals were invoked when participants were explicitly requested to alternately make decisions for themselves or for others. Crucially in the present study, because direct learning and social learning coexisted in the probabilistic environment, and no overt instruction was given to track oneself and the others differently, we argue that these two forms of learning processes are implemented in parallel, and our winning model indicates that the extent to which individuals rely on their own and the others is effectively comparable. Thus, the neurocomputational mechanisms being revealed here are very distinct from those that have been reported previously. Taken collectively, these results demonstrate coexisting, yet distinct value computations in the vmPFC and the ACC for direct learning and social learning, respectively, and are in support of the social-valuation-specific schema^30^.

Our functional connectivity analyses revealed that the mPFC covaried with activations in both vmPFC and ACC. According to a recent meta-analysis^9^, this region is particularly engaged during the decision stage when individuals are representing options and selecting actions, especially in value-based and goal-directed decision-making^40^. Hence, it suggests that beyond the dissociable neural underpinnings, the direct value and vicarious value are further combined to make subsequent decisions^41^.

Furthermore, we replicated previous evidence that NAcc is associated with the RPE computation instead of mere outcome valence^15, 32^. That is, if a brain region encodes the RPE, its activity should be positively correlated with the actual outcome, and negatively correlated with the expected outcome. Beyond reassuring the RPE signal encoded in the NAcc, the corresponding time series analysis serves as a verification framework for testing neural correlates of any error-like signals. As such, our connectivity results seeded at the rTPJ identified a hitherto uncharacterized social prediction error, the difference between actual and expected social outcome, that is encoded in a section of the putamen. This suggests that the SPE signal may trigger a re-computation of expected values and give rise to the subsequent behavioral adjustment. We nonetheless acknowledge that the connectivity analyses here assess correlation rather than directionality, and establishing the casual account by using brain stimulations^42^ or pharmacological manipulations^43^ would be a promising avenue for future work. Albeit this methodological consideration, these functional connectivity results concur with previous evidence that the rTPJ has functional links with the brain’s reward network, of which the striatal region is a central hub^44^.

It is perhaps surprising and interesting that we did not find significant neural correlates with post-decision confidence (i.e., “bet”). This might be due to the fact that decision cues in our current design (i.e., Choice 1, Bet 1, Choice 2, Bet 2) were not presented far apart in time, such that even carefully specified GLMs were not able to capture the variance related to the bets. More importantly, bets in the current design were closely tied to the corresponding choice valuation. In other words, when individuals were sure that one option would lead to a reward, they tended to place a high bet. In fact, this relationship was well-reflected in our winning model and related model parameters (Fig. 2g): bet magnitude was positively correlated with value signals, thus inevitably resulting in co-linear regressors and diminishing the statistical power when assessing its neural correlates. These caveats aside, our results nonetheless shed light on the change in confidence after incorporating social information in decision-making, which goes beyond evidence from previous studies that neither directly addressed the difference in confidence before and after exposing the social information, nor examined the interface between choice and confidence^22, 23^.

It is worth also noting that the model space in the current study is not exhaustive. In particular, we did not test Bayesian models that would track more complex task dynamics^45, 46^, as this class of models may not give advantage in our task environment^47^. The complexity of our task structure, with making four sets of choices and bets and observing two sets of actions as well as the action-outcome associations from four other co-players, made the construction of explicit representation prescribed by Bayesian models rather challenging. In addition, it is so far still unanswered whether RL-like models or Bayesian models provide a more veridical description of how humans make decisions under uncertainty^48^. Regardless of this debate, our fictitious RL model implemented for direct learning is reconciled with previous findings showing its success in reversal learning tasks in both humans^25^ and non-human primates^19^.

In summary, our results provide behavioral, computational, and neural evidence for dissociable representations of direct valuation learned from own experience and vicarious valuation learn from observations of social partners. Moreover, these findings suggest a network of distinct, yet interacting brain regions substantiating crucial computational variables that underlie these two forms of learning. Such a network is in a prime position to process decisions of the sorts mentioned in the beginning, where—as in the example of a lunch order—we have to balance our own experienced-based reward expectations with the expectations of congruency with others and use the resulting error signals to flexibly adapt our choice behavior in social contexts.

## METHODS

### Participants

Forty-one groups of five healthy, right-handed participants were invited to participate in the main study. No one had any history of neurological and psychiatric diseases, nor current medication except contraceptives or any MR-incompatible foreign object in the body. To avoid gender bias, each group consisted of only same-gender participants. To avoid familiarity bias, we explicitly specified in the recruitment that if friends were signing up, they should sign up for different sessions. Forty-one out of 205 participants (i.e., one of each group) were scanned with fMRI while undergoing the experimental task. The remaining 164 participants were engaged in the same task via intranet connections while being seated in the adjacent behavioral testing room outside the scanner. Twenty participants out of 205 who had only switched once or had no switch at all were excluded, including two fMRI participants. This was to ensure that the analysis was not biased by these non-responders (Tomlin et al., 2013). The final sample consisted of 185 participants (95 females; mean age: 25.56 ± 3.98 years; age range: 18-37 years), and among them, 39 participants belonged to the fMRI group (20 females; mean age: 25.59 ± 3.51 years; age range: 20-37 years).

In addition, thirty-nine healthy, right-handed participants were invited to participate in the non-social control study. No one had any history of neurological and psychiatric diseases, nor current medication except contraceptives. To avoid familiarity bias, we explicitly specified in the recruitment that if friends were signing up, they should sign up for different sessions. Extra care during recruitment was taken to exclude participants who had participated in our main study.

Three participants out of 39 who had only switched once or had no switch at all were excluded. This was to ensure that the analysis was not biased by these non-responders^49^. The final sample consisted of 36 participants (19 females; mean age: 23.61 ± 3.42 years; age range: 19-34 years).

All participants in both studies gave informed written consent before the experiment. The study was conducted in accordance with the Declaration of Helsinki and was approved by the Ethics Committee of the Medical Association of Hamburg (PV3661).

### Task design

#### Underlying probabilistic reversal learning paradigm

The core of our social influence task was a probabilistic reversal learning (PRL) task. In our two-alternative forced choice PRL (Supplementary Fig. 1b), each choice option was associated with a particular reward probability (i.e., 70% and 30%). After a variable length of trials (length randomly sampled from a Uniform distribution between 8 and 12 trials), the reward contingencies reversed, such that individuals who were undergoing this task needed to re-adapt to the new reward contingencies so as to maximize their outcome. Given that there was always a “correct” option, which led to more reward than punishment, alongside an “incorrect” option, which caused otherwise, a higher-order anticorrelation structure thus existed to represent the underlying reward dynamics. Such task specification also laid the foundation for our use of fictitious reinforcement learning model with counterfactual updating^25, 50^, .

We used the PRL task rather than tasks with constant reward probability (e.g., always 70%) because the PRL task structure required participants to continuously pay attention to the reward contingency, in order to adapt to the potentially new state of the reward structure and to ignore the (rare) probabilistic punishment from the “correct” option. As a result, the PRL task assured constant learning throughout the entire experiment: choice accuracy reduced after reversal took place, but soon re-established (Supplementary Fig. 2b,c). In fact, one of our early pilot studies used a fixed reward probability. There, participants quickly learned the reward contingency and neglected the social information; thus in this set-up, we could not tease apart the contributions between reward-based influence and socially-based influence.

#### Breakdown of the social influence task (main study)

For each experimental session, a group of five participants were presented with and engaged in the same PRL task via an intranet connection without experimental deception. For a certain participant, portrait photos of the other four same-gender co-players were always displayed within trials (Fig. 1a). This manipulation further increased the ecological validity of the task, at the same time created a more engaging situation for the participants.

The social influence task contained six phases. *Phase 1. Initial choice (Choice 1)*. Upon the presentation of two choice options using abstract fractals, participants were asked to make their initial choice. A yellow frame was then presented to highlight the chosen option. *Phase 2. Initial bet (Bet 1)*. After making Choice 1, participants were asked to indicate how confident they were in their choice, being “1” (not confident), “2” (reasonably confident) or “3” (very confident). Notably, the confidence ratings also served as post-decision wagering metric (an incentivized confidence rating^24, 51, 52^); namely, the ratings would be multiplied by their potential outcome on each trial. For instance, if a participant won on a particular trial, the reward unit (i.e., 20 cent in the current setting) was then multiplied with the rating (e.g., a bet of “2”) to obtain the final outcome (20 × 2 = 40 cent). Therefore, the confidence rating in the current paradigm was referred to as “bet”. A yellow frame was presented to highlight the chosen bet. *Phase 3. Preference giving*. Once all participants had provided their Choice 1 and Bet 1, the choices (but not the bets) of the other co-players were revealed. Crucially, instead of seeing all four other choices at the same time, participants had the opportunity to sequentially uncover their peer’s decisions. In particular, participants could decide whom to uncover first and whom to uncover second, depending on their preference. Choices belonged to the preferred co-players were then displayed underneath the corresponding photo. The remaining two choices were displayed automatically afterward. This manipulation was essential, because, in studies of decision-making, individuals tend to assign different credibility to their social peers based on their performance^15, 21^. And the resulting social preference may play an important role in social decision-making^30^. In the current study, because there were four other co-players in the same learning environment, it was likely that they had various performance levels, and therefore would receive difference preference from the observer. *Phase 4. Choice adjustment (Choice 2)*. When all four other choices were presented, participants were able to adjust their choices given the instantaneous social information. The yellow frame was shifted accordingly to highlight the adjusted choice. *Phase 5. Bet adjustment (Bet 2)*. After the choice adjustment, participants might adjust their bet as well. Additionally, participants also observed other co-players’ Choice 2 (on top of their Choice 1) once they had submitted their adjusted bets. Presenting other co-players’ choices after participants’ bet adjustment rather than their choice adjustment prevented a biased bet adjustment by the additional social information. The yellow frame was shifted accordingly to highlight the adjusted bet. *Phase 6. Outcome delivery*. Finally, the outcome was determined by the combination of participants’ Choice 2 and Bet 2 (e.g., 20 × 2 = 40 cent). Outcomes of the other co-players were also displayed, but shown only as the single reward unit (i.e., 20 cent gain or loss) without being multiplied with their Bet 2. This was to provide participants with sufficient yet not overwhelming information about their peer’s performance. On each trial, the reward was assigned to only one choice option given the reward probability; that is, only choosing one option would lead to a reward, whereas choosing the other option would lead to a punishment. The reward realization sequence (trial-by-trial complementary win and loss) was generated with a pseudo-random order according to the reward probability before the experiment, and this sequence was identical within each group.

### Experimental procedure

To ensure a complete understanding of the task procedure, this study was composed of a two-day procedure: *pre-scanning training* (Day 1), and *main experiment* (Day 2).

#### *Pre-scanning training* (Day 1)

One to two days prior to the MRI scanning, five participants came to the behavioral lab to participate in the pre-scanning training. Upon arrival, they received the written task instruction and the consent form. After returning the written consent, participants were taken through a step-by-step task instruction by the experimenter. Notably, participants were explicitly informed (a) that an intranet connection was established so that they would observe real responses from the others, (b) what probabilistic reward meant by receiving examples, (c) that there was neither cooperation nor competition in this experiment, and (d) that the reward probability could reverse multiple times over the course of the experiment, but participants were not informed about when and how often this reversal would take place. Importantly, to shift the focus of the study away from social influence, we stressed the experiment as a multi-player decision game, where the goal was to detect the “good option” so as to maximize their personal payoff in the end. Given this uncertainty, participants were instructed that they may either trust their own learning experience through trial-and-error, or take decisions from their peers into consideration, as some of them might learn faster than the others. Participants’ explicit awareness of all possible alternatives was crucial for the implementation of our social influence task. To further enhance participants’ motivation, we informed them that the amount they would gain from the experiment would be added to their base payment (see *Reward Payment* below). After participants had fully understood the task, we took portrait photos of them. To avoid emotional arousal, we asked participants to maintain a neutral facial expression as in typical passport photos. To prevent potential confusion before the training task, we further informed participants that they would only see photos of the other four co-players without seeing themselves.

The training task contained 10 trials and differed from the main experiment in two aspects. Firstly, it used a different set of stimuli than those used in the main experiment to avoid any learning effect. Secondly, participants were given a longer response window to fully understand every step of the task. Specifically, each trial began with the stimuli presentation of two choice alternatives and participants were asked to decide on their Choice 1 (4000 ms) and Bet 1 (3000 ms). After the two sequential preference ratings (3000 ms each), all Choice 1 from the other four co-players were displayed underneath their corresponding photos (3000 ms). Participants were then asked to adjust their choice (Choice 2; 4000 ms) and their bet (Bet 2; 3000 ms). Finally, outcomes of all participants were released (3000 ms), followed by a jittered inter-trial interval (ITI, 2000–4000 ms). To help participants familiarize themselves, we orally instructed them what to expect and what to do on each phase for the first two to three trials. The procedure during Day 1 lasted about one hour.

#### *Main experiment* (Day 2)

On the testing day, the five participants came to the MRI building. After a recap of all the important aspects of the task instruction, the MRI participant gave the MRI consent and entered the scanner to perform the main social influence task, while the remaining four participants were seated in the same room adjacent to the scanner to perform the task. All computers were interconnected via the intranet connection. They were further instructed not to make any verbal or gestural communications with other participants during the experiment.

The main experiment consisted of 100 trials and used a different pair of stimuli from the training task. It followed the exact description detailed above (see *Breakdown of the social influence task*; Fig. 1A). Specifically, each trial began with the stimuli presentation of two choice alternatives and participants were asked to decide on their Choice 1 (2500 ms) and Bet 1 (2000 ms). After the two sequential preference ratings (2000 ms each), all Choice 1 from the other four co-players were displayed underneath their corresponding photos (3000 ms). Participants were then asked to adjust their choice (Choice 2; 3000 ms) and their bet (Bet 2; 2000 ms). Finally, outcomes of all participants were released (3000 ms), followed by a jittered inter-trial interval (ITI, 2000–4000 ms). Note that the reward realization sequence (trial-by-trial complementary win and loss) was generated with a pseudo-random order according to the reward probability before the experiment, and this sequence was identical within each group. The procedure during Day 2 lasted about 1.5 hours.

#### Reward payment

All participants were compensated with a base payment of 35 Euro plus the reward they had achieved during the main experiment. In the main experiment, to prevent participants from careless responses on their Choice 1, they were explicitly instructed that on each trial, either their Choice 1 or their Choice 2 would be used to determine the final payoff. However, this did not affect the outcome delivery on the screen. Namely, although on some trials participants’ Choice 1 was used to determine their payment, only outcomes that corresponded to their Choice 2 appeared on the screen. Additionally, when their total outcome was negative, no money was deducted from their final payment. Overall, participants gained 4.48 ± 4.41 Euro after completing the experiment. Finally, the experiment ended with an informal debriefing session.

### Behavioral data acquisition

Stimulus presentation, MRI pulse triggering, and response recording were accomplished with Matlab R2014b (www.mathworks.com) and Cogent2000 (www.vislab.ucl.ac.uk/cogent.php). In the behavioral group (as well as during the pre-scanning training), buttons “V” and “B” on the keyboard corresponded to the left and right choice options, respectively; and buttons “V”, “B”, and “N” corresponded to the bets “1”, “2”, and “3”, respectively. As for the MRI group, a four-button MRI-compatible button box with a horizontal button arrangement was used to record behavioral responses. Buttons “a” and “b” on the button box corresponded to the left and right choice options, respectively; and “a”, “b”, and “c” corresponded to the bets “1”, “2”, and “3”, respectively. To avoid motor artifacts, the position of the two choices options was counterbalanced for all participants.

### MRI data acquisition and pre-processing

MRI data collection was conducted on a Siemens Trio 3T scanner (Siemens, Erlangen, Germany) with a 32-channel head coil. Each brain volume consisted of 42 axial slices (voxel size, 2 × 2 × 2 mm, with 1 mm spacing between slices) acquired using a T2*-weighted echoplanar imaging (EPI) protocol (TR, 2510 ms; TE, 25 ms; flip angle, 40°; FOV, 216 mm) in descending order. Orientation of the slice was tilted at 30° to the anterior commissure-posterior commissure (AC-PC) axis to improve signal quality in the orbitofrontal cortex^53^. Data for each participant were collected in three runs with total volumes ranging from 1210 to 1230, and the first 3 volumes of each run were discarded to obtain a steady-state magnetization. In addition, a gradient echo field map was acquired before EPI scanning to measure the magnetic field inhomogeneity (TE1 = 5.00 ms, TE2 = 7.46 ms), and a high-resolution anatomical image (voxel size, 1 × 1 × 1 mm) was acquired after the experiment using a T1-weighted MPRAGE protocol.

fMRI data preprocessing was performed using SPM12 (Statistical Parametric Mapping; Wellcome Trust Center for Neuroimaging, University College London, London, UK). After converting raw DICOM images to NIfTI format, image preprocessing continued with slice timing correction using the middle slice of the volume as the reference. Next, a voxel displacement map (VDM) was calculated from the field map to account for the spatial distortion resulting from the magnetic field inhomogeneity^54–56^. Incorporating this VDM, the EPI images were then corrected for motion and spatial distortions through realignment and unwarping^55^. The participants’ anatomical images were manually checked and corrected for the origin by resetting it to the AC-PC. The EPI images were then coregistered to this origin-corrected anatomical image. The anatomical image was skull stripped and segmented into gray matter, white matter, and CSF, using the “Segment” tool in SPM12. These gray and white matter images were used in the SPM12 DARTEL toolbox to create individual flow fields as well as a group anatomical template^57^. The EPI images were then normalized to the MNI (Montreal Neurological Institute) space using the respective flow fields through the DARTEL toolbox normalization tool. A Gaussian kernel of 6 mm full-width at half-maximum (FWHM) was used to smooth the EPI images.

After the preprocessing, we further identified brain volumes that (a) excessively deviated from the global mean of the blood-oxygen-level-dependent imaging (BOLD) signals (> 1 SD), (b) showed excessive head movement (movement parameter / TR > 0.4), or (c) largely correlated with the movement parameters and the first derivative of the movement parameters (*R*^2^ > 0.95). This procedure was implemented with the “Spike Analyzer” tool (https://github.com/GlascherLab/SpikeAnalyzer) which returned indices of those identified volumes. We then constructed them as additional participant-specific nuisance regressors of no interest across all our first-level analyses. This implementation identified 3.41 ± 4.79% of all volumes. Note that as this procedure was done per participant, the total number of regressors for each participant may differ.

### Behavioral data analysis

We tested for participants’ behavioral adjustment after observing the instantaneous social information (during Phase 3), by assessing their choice switch probability in Phase 4 (how likely participants switched to the opposite option) and bet difference in Phase 5 (Bet 2 magnitude minus Bet 1 magnitude) as a measurement of how choice and confidence were modulated by the social information. Neither group difference (MRI vs. behavioral) nor gender difference (male vs. female) was observed for the choice switch probability (group: *F*_1,914_ = 0.14, *p* = 0.71; gender: *F*_1,914_ = 0.24, *p* = 0.63) and the bet difference (group: *F*_1,914_ = 0.09, *p* = 0.76; gender: *F*_1,914_ = 1.20, *p* = 0.27). Thus, we pulled data altogether to perform all subsequent analyses. Additionally, trials where participants did not give valid responses on either Choice 1 or Bet 1 in time were excluded from the analyses. On average, 7.9 ± 7.3% of the entire trials were excluded.

We first tested how the choice switch probability and the bet difference varied as a function of the direction of the group (with and against, with respect to each participant’s Choice 1) and the consensus of the group (2:2, 3:1, 4:0, view of each participant; Fig. 1c). To this end, we submitted the choice switch probability and the bet difference to an unbalanced 2 (direction) × 3 (consensus) repeated measures linear mixed-effect (LME) model. The unbalance was due to the fact that data in the 2:2 condition could only be used once, and we grouped it into the “against” condition, thus resulting in three consensus levels in the “against” condition and two consensus levels in the “with” condition. Grouping it into the “with” condition did not alter the results. Furthermore, we further tested the bet difference depending on whether participants switched or stayed on their Choice 2, by performing a 3 (group coherence, 2:2, 3:1, 4:0) × 2 (direction, with vs. against) × 2 (choice type, switch vs. stay) repeated measures LMEs. We constructed LME models with different random effect specifications (Supplementary Table 1) and selected the best one for the subsequent statistical analyses (Fig. 1d,e, Supplementary Fig. 2a). We performed similar analyses with data from the non-social control study (Supplementary Fig. 1c,d).

We further tested whether it was beneficial for the participants to adjust their choice and bet after receiving the instantaneous social information; in other words, we assessed whether participants’ switching behavior was elicited by considering social information or driven by purely perceptual mismatch (i.e., being confronted with visually distinct symbols). We reasoned that if participants were considering social information in our task, the accuracy of their Choice 2 was expected to be higher than that of their Choice 1 (i.e., choosing the “good” option more often). By contrast, if participants’ switching behavior was purely driven by perceptual mismatch, a more random pattern ought to be expected, with no difference between the accuracy of Choice 1 and Choice 2. To this end, we assessed the difference in the accuracy between Choice 1 and Choice 2 (Fig. 1f), as well as the difference of the magnitude between Bet 1 and Bet 2 (Fig. 1g), using two-tailed paired *t*-tests. We also tested how choice accuracy and bet magnitude changed across reversals. We selected a window of seven trials (three before and three after reversal, reversal included) to perform this analysis, with data being stacked with respect to the reversal (i.e., trial-locked) and averaged per participants. We submitted the data to a 2 (Choice 1 vs. Choice 2 or Bet 1 vs. Bet 2) × 7 (relative trial position, −3, −2, −1, 0, +1, +2, +3) repeated measures LME models with five different random effect specifications, respectively (Supplementary Table 2). When the main effect of *position* was significant, we submitted the data to a post-hoc comparison with Tukey’s HSD correction (Supplementary Fig. 2b,c). We performed similar analyses with data from the non-social control study (Supplementary Fig. 1e.,f).

In addition, although we did not intentionally manipulate the amount of dissenting social information (given the real-time property of our task), the sequence was nonetheless randomly progressed for nearly all participants (Wald-Wolfowitz runs test showed 178 out of 185 participants’ trial-by-trial amount of dissenting social information was randomly ordered, *p*s > 0.05). In order to guard against possible confounding effects, we nonetheless tested whether the amount of dissenting social information and participants’ behavior was related to task structure (time of reversal) and participants’ lapse error. Note that, the lapse error was defined as choosing one choice option on Choice 1 when the model strongly favored the alternative (modeled action probability >= 95%). For example, when the model predicted *p*(A) of Choice 1 was 95% (or higher) yet the participants actually chose option B, this trial was referred to as a lapse error. We tested the Pearson’s correlation between the following pairs of variables for each participant and for the MRI participants: (a) amount of dissenting social information and time of reversal, (b) amount of dissenting social information and lapse error (c) participants’ switching behavior and time of reversal, and (d) participants’ switching behavior and lapse error. Results indicated no significant relationship between any of the above pairs of variables (Supplementary Fig. 2d,e).

All statistical tests were performed in R (v3.3.1; www.r-project.org). All repeated-measures LME models were analyzed with the “lme4” package^58^ in R. Results were considered statistically significant at the level *p* < 0.05.

### Computational modeling

To describe participants’ learning behavior in our social influence task and to uncover latent trial-by-trial measures of decision variables, we developed three categories of computational models and fitted these models to participants’ behavioral data. We based all our computational models on the simple reinforcement learning model (RL^5^), and progressively include components (Table 1).

First, given the structure of the PRL task, we sought to evaluate whether a fictitious update RL model that incorporates the anticorrelation structure (see *Underlying probabilistic reversal learning paradigm*) outperformed the simple Rescorla-Wagner^28^ RL model that only updated the value of the chosen option and the Pearce-Hall^59^ model that employed a dynamic learning rate to approximate the optimal Bayesian learner. These models served as the baseline and did not consider any social information (Category 1: M1a, M1b, M1c). On top of Category 1 models, we then included the instantaneous social influence (i.e., other co-players’ Choice 1, before outcomes were delivered) to construct social models (Category 2: M2a, M2b, M2c). Finally, we considered the component of social learning with competing hypotheses of value update from observing others (Category 3: M3, M4, M5, M6a, M6b). The remainder of this section explains choice-related model specifications and bet-related model specifications (see Supplementary Table 3 for a list of full specifications).

#### Choice model specifications

In all models, Choice 1 was accounted for by the option values of option A and option B:

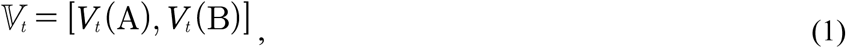

where *V_t_* indicated a two-element vector consisting of option values of A and B on trial *t*. Values were then converted into action probabilities using a Softmax function^5^. On trial *t*, the action probability of choosing option A (between A and B) was defined as follows:

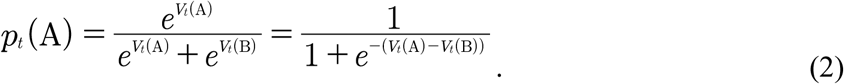

For Choice 2, we modeled it as a “switch” (1) or a “stay” (0) using a logistic regression. On trial *t*, the probability of switching given the switch value was defined as follows:

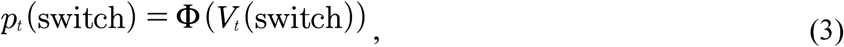

where Φ was the inverse logistic linking function:

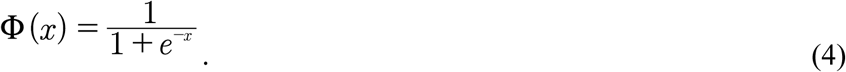

It is worth noting that, in model specifications of the action probability, we did not include the commonly-used inverse Softmax temperature parameter *τ*. This was because we explicitly constructed the option values of Choice 1 and the switch value of Choice 2 in a design-matrix fashion (e.g., Eq. 6; and see the text below). Therefore, including the inverse Softmax temperature parameter would inevitably give rise to a multiplication term, which, as a consequence, would cause unidentifiable parameter estimation^27^. For completeness, we also assessed models with the *τ* parameter, and they performed consistently worse than our models specified here.

The Category 1 models (M1a, M1b, M1c) did not consider any social information. In the simplest model (M1a), a Rescorla-Wagner model was used to model the Choice 1, with only the chosen value being updated via the reward prediction error (RPE; *δ*), and the unchosen value remaining the same as the last trial.

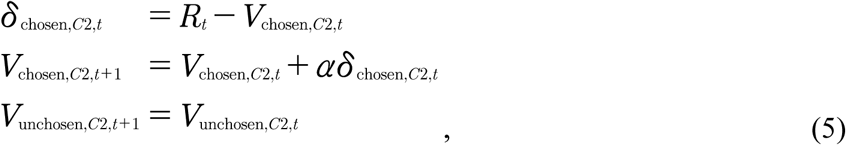

where *R_t_* was the outcome on trial *t*, and *α* (0 < *α* < 1) denoted the learning rate that accounted for the weight of RPE in value update. A beta weight (*β*_V_) was then multiplied by the values before being submitted to Eq. 2 with a Categorical distribution, as in:

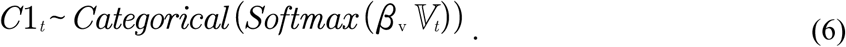

Because there was no social information in M1a, the switch value of Choice 2 was comprised merely of the value difference of Choice 1 and a switching bias (i.e., intercept):

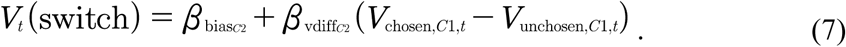

Choice 2 was then modeled with this switch value following a Bernoulli distribution:

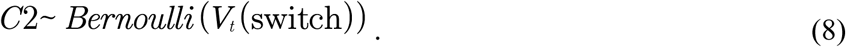

In M1b we tested whether the fictitious update could improve the model performance, as the fictitious update has been successful in PRL tasks in non-social contexts^25, 50^. In M1b, both the chosen value and the unchosen value were updated, as in:

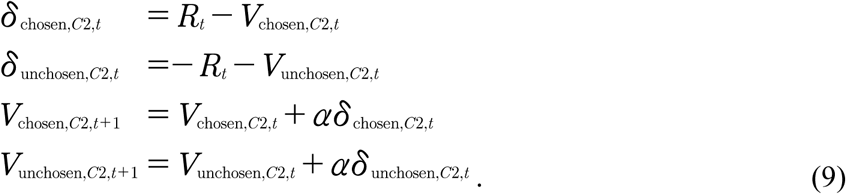

In M1c we assessed the Pearce-Hall^59^ model that entailed a dynamic learning rate, as previous studies have shown its usefulness in associative learning^60^:

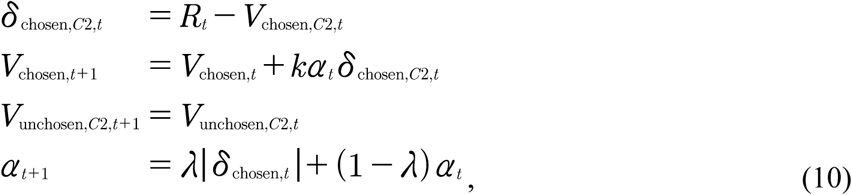

where *k* (0 < *k* < 1) was the weight of the (dynamic) learning rate, and *λ* (0 < *λ* < 1) indicated the weight between RPE and the learning rate.

Our Category 2 models (M2a, M2b, M2c) tested the role of instantaneous social influence on Choice 2, namely, whether observing choices from the other co-players in the same learning environment contributed to the choice switching. As compared with M1 (M1a, M1b, M1c), only the switch value of Choice 2 was modified, as follows:

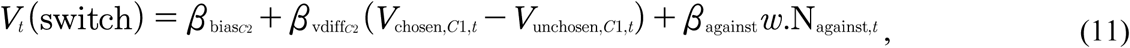

where *w*.N_against,*t*_ denoted the preference-weighted amount of dissenting social information relative to each participant’s Choice 1 on trial *t*. It was computed on a trial-by-trial fashion as follows:

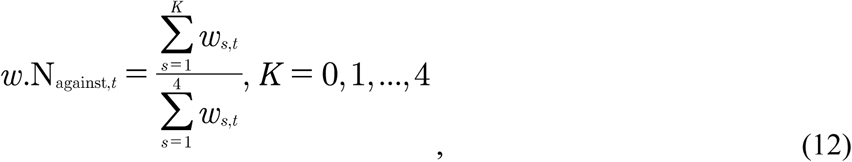

where *K* indicated the number of opposite choices from the others, *w_s_*_,*t*_ was participants’ trial-by-trial preference weight toward the other four co-players. Note that these preference weights were fixed parameters based on each participant’s preference toward the others when uncovering their choices: the 1^st^ favored co-player received a weight of 0.75, the 2^nd^ favored co-player received a weight of 0.5, and the rest two co-players received a weight of 0.25, respectively. They were not modeled as free parameters because doing so caused unidentifiable model estimate behavior. All other specifications of models in this category (M2a, M2b, M2c) were identical to models in Category 1 (M1a, M1b, M1c), respectively.

Our Category 3 models (M3, M4, M5, M6a, M6b) assessed whether participants learned from their social partners and whether they updated vicarious option values through social learning. It is worth noting that, models belonging to Category 2 solely considered the instantaneous social influence on Choice 2, whereas models in Category 3 tested several competing hypotheses of the vicarious valuation that may contribute to Choice 1 on the following trial, in combination with individuals’ own valuation processes. In all models within this category, the option values of Choice 1 was specified by a weighted combination between *V*_self_ updated via direct learning and *V*_other_ updated via social learning:

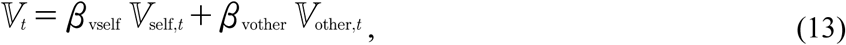

Where

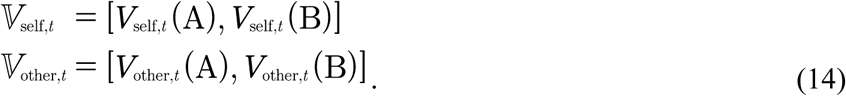

Note that given M2b was the winning model among Category 1 and Category 2 models (Table 1), we used M2b’s specification for the value update of *V*_self_ (Eq. 9), so that Category 3 models only differed on the specification of *V*_other_.

M3 tested whether individuals recruited a similar RL algorithm to their own when learning option values from observing others. As such, M3 assumed participants to update values “for” the others using the same fictitious update rule for themselves (Eq. 7):

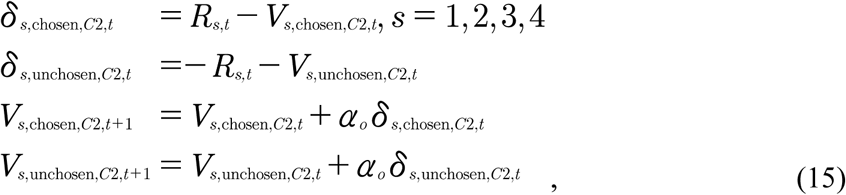

where *s* denoted the index of the four other co-players. These option values from the four co-players were then preference-weighted and summed to formulate *V*_other_, as follows:

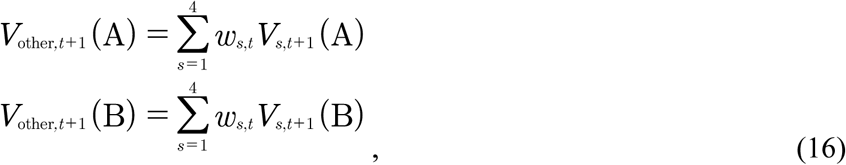

where *w_s_*_,*t*_ was participants’ preference weight. To ensure that the corresponding value-related parameters (*β*_vself_ and *β*_vother_ in Eq. 13) were comparable, *V*_other_ was further normalized to lie between −1 and 1 with the Φ(x) function defined in Eq. 4:

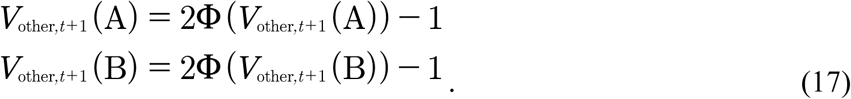

One may argue that having four independent RL agents as in M3 was cognitively demanding: in order to accomplish so, participants had to track and update each other’s individual learning processes together with their own valuation (together 2^5^ units of information). We, therefore, constructed three additional models that employed simpler but distinct pathways to update vicarious values via social learning. In essence, M3 considered both choice and outcome to determine the action value. We then asked if using either choice or outcome alone may perform as well as, or even better than, M3. Following this assumption, we constructed (a) M4 that updated *V*_other_ using only the others’ action preference, (b) M5 that considered the others’ current outcome to resemble the value update via observational learning, and (c) M6a that tracked the others’ cumulative outcome to resemble the value update via observational learning.

In M4, other players’ action preference (*ρ*) is derived from the choice history over the last three trials using the cumulative distribution function of the beta distribution at the value of 0.5 (*I*_0.5_). That is:

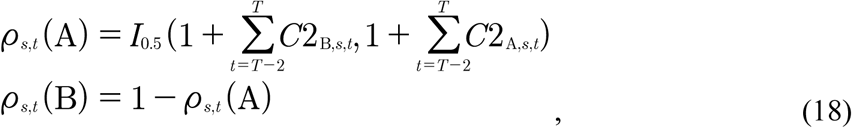

where *s* denoted the index of the four other co-players, *t* denoted the trial index from *T*−2 to *T*. To illustrate, if one co-player chose option A twice and option B once in the last three trials, then the action preference of choosing A for him/her was: *I*_0.5_(frequency of B + 1, frequency of A + 1) = *I*_0.5_(0.5, 1 + 1, 2 + 1) = 0.6875. *V*_other_ was computed based on these action preferences:

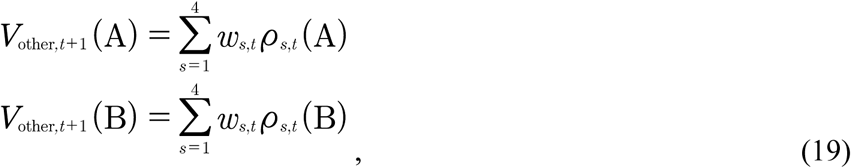

where *w_s_*_,*t*_ was participants’ preference weight, and *s* denoted the index of the four other co-players. Like M3, the computation of *V*_other_ here was also preference-weighted and summed. The values were similarly normalized using Eq. 17.

By contrast, M5 tested whether participants updated *V*_other_ using only each other’s reward on the current trial:

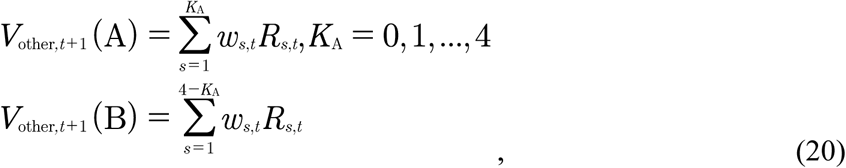

where *w_s_*_,*t*_ was participants’ preference weight, *s* denoted the index of the four other co-players, *t* denoted the trial index from *T*−2 to *T*, and *K*_A_ denoted the number of co-players who decided on option A on trial *t*. Like M3, the computation of *V*_other_ here was also preference-weighted and summed. These values were then normalized using Eq. 17.

Moreover, M6a assessed whether participants tracked a cumulated reward histories over the last few trials instead of monitoring only the most recent outcome of the others. In fact, a discounted reward history over the recent past (e.g., the last three trials) has been a relatively common implementation in other RL studies in non-social contexts^29, 61, 62^. By testing four window sizes of trials (i.e., three, four, or five) and using a nested model comparison, we decided on a window of three past trials to accumulate the other co-players’ performance:

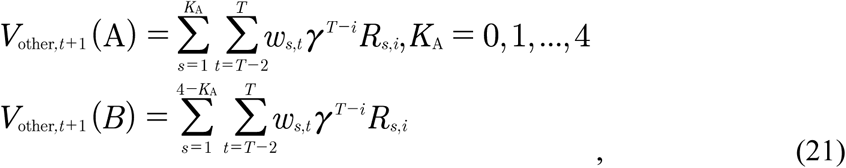

where *γ* (0 < *γ* < 1) denoted the rate of exponential decay, all other notions were as in Eq. 20. Like M3, the computation of *V*_other_ here was also preference-weighted and summed. The values were then normalized using Eq. 17.

Lastly, given that M6a was the winning model among all the models above (M1 – M6a) indicated by model comparison (see below Model selection; Table S1), we further assessed in M6b whether Bet 1 contributed to the choice switching on Choice 2, as follows:

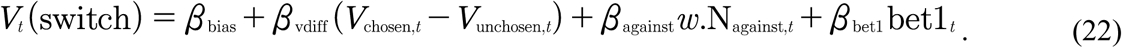

It is noteworthy that in M6a/M6b, *V*_other_ differed from *V*_self_ in practice. On trial *t*, *V*_self_ of a punished option might largely decrease given the negative RPE, whereas *V*_other_ may not be vastly affected because of the others’ previous successes (e.g., *V*_other_(Blue) Fig. 2c: albeit a loss on trial *t*, the cumulative reward history was still positive, indicating the cumulative performance was still reliable). In fact, both *V*_self_ and *V*_other_ spanned within their range (−1 to 1; Fig. 2d) with a slightly moderate correlation (*r* = 0.38 ± 0.097 across participants; Fig. 3a), and they jointly contributed to the action probability of Choice 1.

#### Bet model specificationsthe winning model

In all models, both Bet 1 and Bet 2 were modeled as ordered-logistic regressions that are often used for quantifying discrete variables, like Likert-scale questionnaire data^63^. We applied the ordered-logistic model because the bets in our study indeed inferred an ordinal feature. Namely, betting on three was higher than betting on two, and betting on two was higher than betting on one, but the difference between the bets of three and one (i.e., a difference of two) was not necessarily twice as the difference between the bets of three and two (i.e., a difference of one). Hence, we sought to model the distance (decision boundary) between them. Moreover, we hypothesized a continuous computation process of bet utilities when individuals were placing bets, which satisfied the general assumption of the ordered-logistic regression model.

There were two key components in our bet models, the continuous bet utility *U*_bet_, and the set of boundary thresholds *θ*. Specifically, the bet utility *U*_bet_ varied between *K*−1 thresholds (*θ*_1, 2, …, *K*-1_) thresholds to predict bets. Since there were three bet levels in our task (*K* = 3), we introduced two decision thresholds, *θ*_1_ and *θ*_2,_ (*θ*_2_ > *θ*_1_). As such, the predicted bets (bêt) on trial *t* were represented as follows:

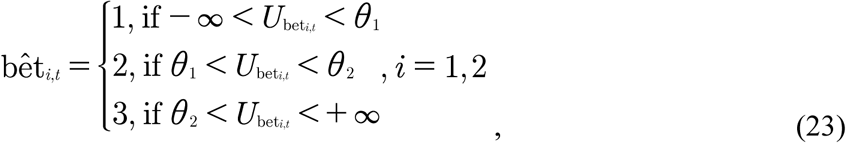

where *i* indicated either bêt 1 or the bêt 2. Because there were only two levels of threshold, for simplicity, we set *θ*_1_ = 0, and *θ*_2_ = *θ,* (*θ* > 0). To model the actual bets, a logistic function (Eq. 4) was used to obtain the action probability of each bet, as follows:

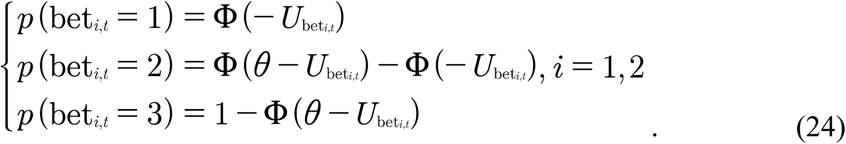

The utility *U*_bet1_ was comprised of a bet bias and the value difference between the chosen option and the unchosen option:

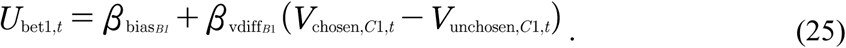

The rationale was that the larger the value difference between the chosen and the unchosen options, the more confident individuals were expected to be, hence placing a higher bet. This utility *U*_bet1_ was kept identical across all models (M1a – M6b), and Bet 1 was modeled as follows:

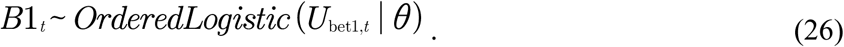

In addition, Bet 2 was modeled as the bet change relative to Bet 1. Therefore, the utility *U*_bet2_ was constructed on top of *U*_bet1_. In all non-social models (M1a, M1b, M1c), the bet change term was represented by a bet change bias (i.e., intercept), depending on whether participants had a switch or stay on their Choice 2:

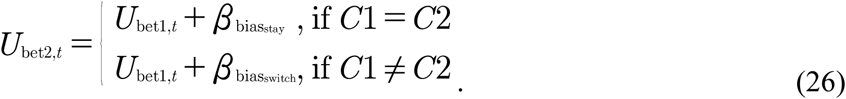

In all social models (M2a – M6b), regardless of the observational learning effect, the bet change term was specified by the instantaneous social information together with the bias, depending on whether participants had a switch or stay on their Choice 2:

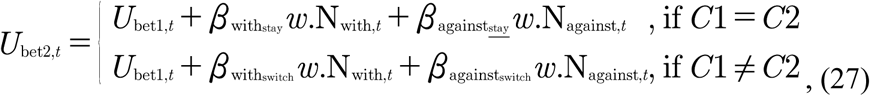

Where

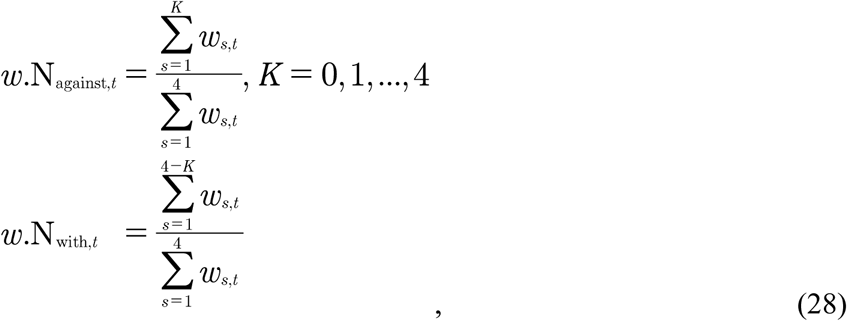

where *K* indicated the number of opposite choices from the others, *w_s_*_,*t*_ was participants’ trial-by-trial preference weight toward the other four co-players. It should be noted that, however, despite the high negative correlation between *w*.N_with_ and *w*.N_against_, the parameter estimation results showed that the corresponding effects (i.e., *β*_with_ and *β*_against_) did not rely on each other (*r* = 0.04, *p* > 0.05). In fact, as shown in Fig. 2H, the corresponding parameters showed independent contributions to the bet change during the adjustment. Additionally, we constructed two other models using either *w*.N_with_ or *w*.N_against_ along, but both model performance dramatically reduced than including both of them (ΔLOOIC > 1000). Lastly, the utility *U*_bet2_ was kept identical across all social models (M2a – M6b), and Bet 2 was modeled as follows:

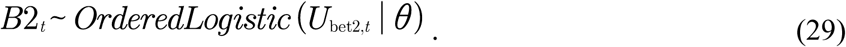

### Hierarchical Bayesian model estimation

In all models, we simultaneously estimated both choices (Choice 1, Choice 2) and bets (Bet 1, Bet 2). Model estimations of all aforementioned candidate models were performed with hierarchical Bayesian analysis^27^ (HBA) using a newly developed statistical computing language Stan^64^ in R. Stan utilizes a Hamiltonian Monte Carlo (HMC; and efficient Markov Chain Monte Carlo, MCMC) sampling scheme to perform full Bayesian inference and obtain the actual posterior distribution. We performed HBA rather than maximum likelihood estimation (MLE) because HBA provides much more stable and accurate estimates than MLE^27^. Following the approach in the “hBayesDM” package^65^ for using Stan in the field of reinforcement learning, we assumed, for instance, that a generic individual-level parameter *φ* was drawn from a group-level normal distribution, namely, *φ* ∼ Normal (*μφ*, *σφ*), with *μφ* and *σφ*. being the group-level mean and standard deviation, respectively. Both these group-level parameters were specified with weakly-informative priors^27^: *μφ* ∼ Normal (0, 1) and *σφ*.∼ half-Cauchy (0, 5). This was to ensure that the MCMC sampler traveled over a sufficiently wide range to sample the entire parameter space. All parameters were unconstrained except for *α* and *γ* (both [0 1] constraint, with inverse probit transform) and *θ* (positive constraint, with exponential transform).

In HBA, all group-level parameters and individual-level parameters were simultaneously estimated through the Bayes’ rule by incorporating behavioral data. We fit each candidate model with four independent MCMC chains using 1000 iterations after 1000 iterations for the initial algorithm warmup per chain, which resulted in 4000 valid posterior samples. The convergence of the MCMC chains was assessed both visually (from the trace plot) and through the Gelman-Rubin 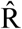 Statistics^66^. 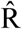 values of all parameters were close to 1.0 (at most smaller than 1.1 in the current study), which indicated adequate convergence.

### Model selection and posterior predictive check

For model comparison and model selection, we computed the Leave-One-Out information criterion (LOOIC) score per candidate model^67^. The LOOIC score provides the point-wise estimate (using the entire posterior distribution) of out-of-sample predictive accuracy in a fully Bayesian way, which is more reliable compared to point-estimate information criterion (e.g., Akaike information criterion, AIC; deviance information criterion, DIC). By convention, lower LOOIC score indicates better out-of-sample prediction accuracy of the candidate model. Plus, a difference score of 10 on the information criterion scale was considered decisive^68^. We selected the model with the lowest LOOIC as the winning model. We additionally performed Bayesian model averaging (BMA) with Bayesian bootstrap^69^ to compute the probability of each candidate model being the best model. Conventionally, BMA probability of 0.9 (or higher) is a decisive indication.

Moreover, given that model comparison provided merely relative performance among candidate models^70^, we then tested how well our winning model’s posterior prediction was able to replicate the key features of the observed data (a.k.a., posterior predictive checks, PPCs). To this end, we applied a post-hoc absolute-fit approach^71^ that factored in participants’ actual action and outcome sequences to generate predictions with the entire posterior MCMC samples. Namely, we let the model generate choices and bets as many times as the number of samples (i.e., 4000 times) per trial per participants and we analyzed the generated data the same way as we did for the observed data, and assessed whether these analyses could reproduce the behavioral pattern in our behavioral analysis (Fig. 1d,e, Supplementary Fig. 2a).

Lastly, we tested how specific model parameters linked with behavioral findings to assess individual differences (Fig. 2i–j). In the choice model, we tested the Pearson’s correlation between *β*(*w*.N_against_) and the first-order polynomial slope derived from the choice switch probability as a function of the group consensus in the “against” condition (as in Fig. 1d, red line). Likewise, in the bet model, we tested the Pearson’s correlation between *β*(*w*.N_with_) and the first-order polynomial slope derived from the bet difference as a function of the group consensus in the “with” condition (as in Fig. 1e, blue line).

### MRI data analysis

#### Deriving internal computational signals

Based on the winning model (Table 1) and its parameter estimation (Fig. 2e–h), we derived trial-by-trial computational signals for each individual MRI participant using the mean of the posterior distribution of the parameters. We used the mean rather than the mode (i.e., the peak resulted from kernel density estimate) because in MCMC, especially HMC implemented in Stan, the mean is much more stable than the mode to serve as the point estimate of the entire posterior distribution^64^. In fact, as we modeled all parameters as normal distributions, the posterior mean and the posterior mode were highly correlated (*r* = 0.99, *p* < 1.0 × 10^−10^). For each MRI participant, we derived the following trial-by-trial variables and behaviors: *V*_self_, *V*_other_, *w*.N_against_, Choice 2 behavior (SwSt: switch vs. stay), *U*_bet1_, *U*_bet2_, RPE.

#### First-level analysis

fMRI data analyses were performed using SPM12. We conducted model-based fMRI analyses^25, 31^ containing the computational signals described above. We set up two event-related general linear models (GLM 1 and GLM 2) to test the neural correlates of decision variables.

GLM 1 assessed the neural representations of valuation resulted from participants’ direct learning and observational learning in Phase 1, as well we the instantaneous social influence in Phase 3. The first-level design matrix in GLM 1 consisted of constant terms, nuisance regressors identified by the “Spike Analyzer”, plus the following 22 regressors: five experimentally measured onset regressors for each cue (cue of Choice 1: 0 s after trial began; cue of Bet 1: 2.92 s after trial began; cue of Choice 2: 12.82 s after trial began; cue of Bet 2: 16.25 s after trial began; cue of outcome: 21.71 s after trial began); six parametric modulators (PM) of each corresponding cue (*V*_self,chosen_, *V*_other,chosen_, belonging to the cue of Choice 1; *w*.N_against_ belonging to the cue of Choice 2; *U*_bet1_, *U*_bet2_, belonging to the cue of Bet 1 and Bet 2, respectively; and RPE belonging to the cue of outcome); five nuisance regressors accounted for all of the “no response” trials (missing trials) of each cue; and six movement parameters. Note that though the two value (*V*_self,chosen_, *V*_other,chosen_) signals were slightly moderately correlated (*r* = 0.38 ± 0.097 across participants; Fig. 3A), *V*_other,chosen_ was orthogonalized with respect to *V*_self,chosen_. This allowed us to obtain as much variance as possible on the *V*_self,chosen_ regressor, and then any additional (explainable) variance would be accounted for by the *V*_other,chosen_ regressor^72^. Also, we intentionally did not include the actual reward outcome at the outcome cue. This was because (a) the RPE and the reward outcome are known to be correlated in goal-directed learning studies using model-based fMRI^71^, and (b) we sought to explicitly verify RPE signals by its hallmarks using the region of interest (ROI) time series extracted from each participant given the second-level RPE contrast (see below *ROI time series analysis*).

GLM 2 was set up to examine the neural correlates of choice adjustment in Phase 4. To this end, GLM 2 was identical to GLM 1, except that the PM regressor of *w*.N_against_ under the onset cue of Choice 2 was replaced by the PM regressor of SwSt (“switch” = 1, “stay” = −1). Additionally, albeit that we showed no pattern between participants’ behavior and task structure (Supplementary Fig. 2d,e), we included each participants’ time of reversal and their lapse error as covariates in GLM 1 and GLM 2, as GLM 3 and GLM 4. Given the non-correlation between variables of interest and the task structure, significant clusters resulted from GLM 3 and GLM 4 nearly identical with those from GLM 1 and GLM 2, respectively.

#### Second-level analysis

The resulting *β* images from each participant’s first-level GLMs were then used in random-effects group analyses at the second level, using one-sample two-tailed *t*-tests for significant effects across participants. To correct for multiple comparisons of functional imaging data, we employed the threshold-free cluster enhancement (TFCE^73^) implemented in the TFCE Toolbox (dbm.neuro.uni-jena.de/tfce/). TFCE is a cluster-based thresholding method that aims to overcome the shortcomings of choosing an arbitrary cluster size (e.g., *p* < 0.001, cluster size *k* = 20) to form a threshold. The TFCE procedure took the raw statistics from the second-level analyses and performed a permutation-based non-parametric test (i.e., 5000 permutations in the current study) to obtain robust results. Plus, given our hypotheses and according to existing evidence that vmPFC encodes experiential value signals from direct learning^9^ and that ACC tracks vicarious value signals from social learning^15, 21, 38^, we performed small volume corrections (SVC) for the value related contrast using 10-mm search volumes around the peak MNI coordinates of the vmPFC (x = 2, y = 46, z = −8) and the ACC (x = 2, y = 14, z = 30) reported in the corresponding studies with the TFCE correction at *p* < 0.05 (Fig. 3b). For the otherwise whole-brain analyses, we performed whole-brain TFCE correction at *p* < 0.05, FWE (family-wise error) corrected (Fig. 3c, Supplementary Fig. 4, Supplementary Fig. 5).

#### Follow-up ROI analysis

Depending on the hypotheses, the research question, and the corresponding PM regressors, we employed two types of follow-up ROI analyses: the time series estimates and percent signal change (PSC) estimates. In both types of ROI analyses, participant-specific masks were created from the second-level contrast. We applied a previously reported leave-one-out procedure^25^ to extract cross-validated BOLD time series. This was to provide an independent criterion for ROI identification and thus ensured statistical validity^74^. For each participant, we first defined a 10-mm search volume around the peak coordinate of the second level contrast re-estimated from the remaining N−1 participants (threshold: *p* < 0.001, uncorrected); within this search volume, we then searched for each participant’s nearest individual peak and created a new 10-mm sphere around this individual peak as the ROI mask. Finally, supra-threshold voxels in the new participant-specific ROI were used for both ROI analyses.

The ROI time series estimates were applied when at least two PMs were associated with each ROI. Namely, we were particularly interested in how the time series within a specific ROI correlated with all the PM regressors. In the current studies, we defined 3 ROIs to perform the time series estimates: the vmPFC, the ACC, and the VS/NAcc.

We followed the procedure established by previous studies^15, 32^ to perform the ROI time series estimates. We first extracted raw BOLD time series from the ROIs. The time series of each participant was then time-locked to the beginning of each trial with a duration of 30 s, where the cue of Choice 1 was presented at 0s, the cue of Bet 1 was presented at 2.92 s, the cue of Choice 2 was displayed at 12.82 s, the cue of Bet 2 was displayed at 16.25 s, and the cue of outcome was presented at 21.71 s. All these time points corresponded to the mean onsets for each cue across trials and participants. Afterward, ROI time series were up-sampled to a resolution of 250 ms (1/10 of TR) using 2D cubic spline interpolation, resulting in a data matrix of size *m* × *n*, where *m* was the number of trials, and *n* was the number of the up-sampled time points (i.e., 30 s / 250 ms = 120 time points). A linear regression model containing the PMs was then estimated at each time point (across trials) for each participant. It should be noted that, although the linear regression here took a similar formulation as the first-level GLM, it did not model any specific onset; instead, this regression was fitted at each time point within the entire trial across all trials. The resulting time courses of effect sizes (regression coefficients, or *β* weights) were finally averaged across participants. Because both the time series and the PMs were normalized, these time courses of effect sizes, in fact, reflected the partial correlation between the ROI time series and PMs.

To test the group-level significance of the above ROI time series analysis, we employed a non-parametric permutation procedure. For the time sources of effect sizes (*β* weights) for each ROI, we defined a time window of 3–7 s after the corresponding event onset, during which the BOLD response was expected to peak. In this time window, we randomly flipped the signs of the time courses of *β* weights for 5000 repetitions to generate a null distribution, and assessed whether the mean of the generated data from the permutation procedure was smaller or larger than 97.5% of the mean of the empirical data.

Further, the Percent signal change (PSC) estimates were applied when only one PM was associated with each ROI. Particularly, we tested whether there was a linear trend of the PSC for each ROI as a function of the PM. In the current study, we defined seven ROIs to perform the PSC estimates. Among them, four ROIs were associated with the PM regressor of *w*.N_against_, being the rTPJ, the ACC/pMFC, the right aINS and the FPC; two ROIs were associated with the PM regressor of SwSt, being the left dlPFC and the ACC; and one ROI was associated with the inverse contrast of SwSt (i.e., StSw, stay vs. switch), being the vmPFC.

To compute the PSC, we used the “rfxplot” toolbox^75^ to extract the time series from the above ROIs. The “rfxplot” toolbox further divided the corresponding PMs into different bins (e.g., in the case of two bins, PMs were split into the first 50% and the second 50%) and computed the PSC for each bin, which resulted in a *p* × *q* PSC matrix, where *p* was the number of participants, and *q* was the number of bins. To test for significance, we performed a simple first-order polynomial fit using the PSC as a function of the binned PM, and tested whether the slope of this polynomial fit was significantly different from zero using two-tailed one sample t-tests.

#### Functional connectivity analysis

We conducted two types of functional connectivity analyses^76^ in the current study, the psychophysiological interaction (PPI) and the physiophysiological interaction (PhiPI) to assess the functional network using fMRI. In both types of connectivity analyses, the seed brain regions were determined based on the activations from the earlier GLM analyses, and extract cross-validated BOLD time series from each corresponding ROI using the leave-one-out procedure described above.

The psychophysiological interaction (PPI) analysis aims to uncover how the functional connectivity between BOLD signals in a particular ROI (seed region) and BOLD signals in the (to-be-detected) target region(s) is modulated by a psychological variable. We used as a seed the entire BOLD time series from a 10-mm spherical ROI in the rTPJ, centered at the peak coordinates from the PM contrast of *w*.N_against_ (threshold: *p* < 0.001, uncorrected), which was detected at the onset cue of the second choice. Next, we constructed the interaction regressor of the PPI analysis (i.e., the regressor of main interest) by combining the rTPJ ROI signals with the SwSt (“switch” = 1, “stay” = −1) variable that took place at the onset cue of Choice 2. We first normalized the physiological and psychological terms and then multiplied them together, further orthogonalizing their product to each of the two main effects. These three regressors (i.e., the interaction, the BOLD time series of the seed region, and the modulating psychological variable) were finally mean-corrected and then entered into the first-level PPI design matrix. To avoid possible confounding effects, we further included all the same nuisance regressors as the above first-level GLMs: five nuisance regressors accounted for all the “no response” trials (missing trials) of each event cue, six movement parameters, and additional regressors of interest identified by the “Spike Analyzer”. The resulting first-level interaction regressor from each participant was then submitted to a second-level *t*-test to establish the group-level connectivity results, with whole-brain TFCE correction at *p* < 0.05, FWE corrected (Fig. 4a–c).

The Physiophysiological interaction (PhiPI) analysis follows the same principles as the PPI analysis, except that the psychological variable in the PPI regressors is replaced by the BOLD time series from a second seed ROI. For the interaction term, we first normalized the BOLD time series of the two seed regions, and then multiplied them together, further orthogonalizing their product to each of the two main effects. The three regressors (i.e., two main-effect terms and their interaction) were finally mean-corrected and then entered into the first-level PhiPI design matrix.

We performed two PhiPI analyses. In the first PhiPI, we used as seed regions the entire BOLD time series in two 10-mm spherical ROIs in the vmPFC (seed 1) and the ACC (seed 2), both of which were detected at the cue of Choice 1 from the parametric modulators of *V*_self_ and *V*_other_, respectively. The design matrix of the first PhiPI analysis thus consisted of the interaction term between vmPFC and ACC, and the two main-effect regressors with the BOLD time series of vmPFC and ACC, respectively. In the second PhiPI, we seeded with the entire BOLD time series from an identical 10-mm spherical ROI in the rTPJ (seed 1) as described in the above PPI analysis, and from a 10-mm spherical ROI in the left dlPFC (seed 2), which was identified at the cue of Choice 2 from the contrast of choice adjustment (switch > stay). The design matrix of the second PhiPI analysis thus consisted of the interaction term between rTPJ and left dlPFC, and the two main-effect regressors with the BOLD time series of rTPJ and left dlPFC, respectively. In both PhiPI analyses, we further included all the same nuisance regressors as the above first-level GLMs to avoid possible confounding effects: five nuisance regressors accounted for all the “no response” trials (missing trials) of each event cue, six movement parameters, and additional regressors of interest identified by the “Spike Analyzer”. The resulting first-level interaction regressor from each participant was then submitted to a second-level *t*-test to establish the group-level connectivity results, with whole-brain TFCE correction at *p* < 0.05, FWE corrected (Fig. 4e–i, Supplementary Fig. 6a).

## SUPPLEMENTARY INFORMATION

Supplemental Information includes 6 figures, 3 tables, and 2 notes can be found with this article at xxx.

## ACKNOWLEDGMENTS

We thank Anne Bert, Kiona Weisel, Julia Spilcke-Liss, Julia Majewski, and all radiographers for help with data acquisition; Nathaniel Daw for help in developing the computational models; and Christian Büchel for helpful feedback on earlier versions of the manuscript. L.Z. was supported by the International Research Training Groups “CINACS” (DFG GRK 1247), and the Research Promotion Fund (FFM) for young scientists of the University Medical Center Hamburg-Eppendorf. J.G. was supported by the Bernstein Award for Computational Neuroscience (BMBF 01GQ1006), the Collaborative Research Center “Cross-modal learning” (DFG TRR 169), and the Collaborative Research in Computational Neuroscience (CRCNS) grant (BMBF 01GQ1603).

## AUTHOR CONTRIBUTIONS

J.G. conceived the initial research idea. L.Z. performed behavioral pilot testing. L.Z. and J.G. designed and programmed final experiments. L.Z. acquired data. L.Z. and J.G. designed computational models. L.Z. and J.G. performed analyses, interpreted the results, and wrote the manuscript. J.G. supervised the project.

