## Supplemental Information for "A brain network supporting social influences in human decision-making"

### Supplementary Figures

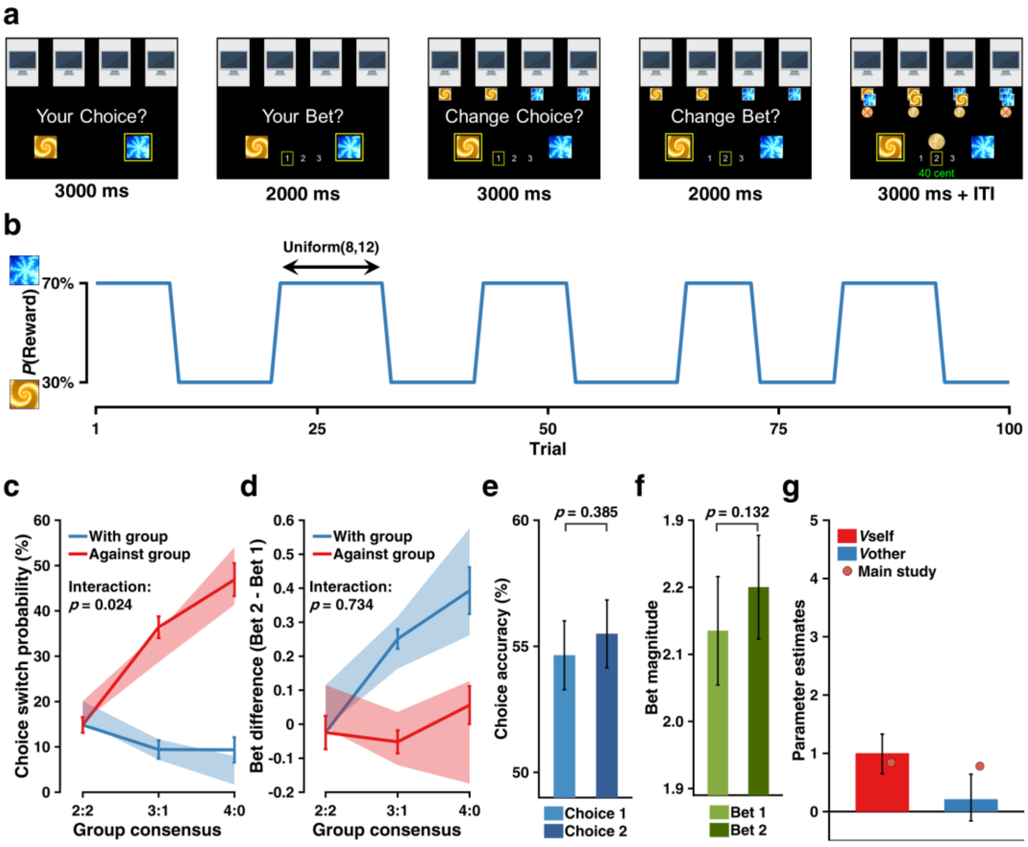


**Supplementary Figure 1**

**Non-social control experiment.**

**(a)** Task design. Participants (N = 36) made an initial choice and an initial bet (Choice 1, Bet 1), and after observing the other four intelligent computer agents' initial choices (Choice 2, Bet 2), they were asked to adjust their choice and bet, followed by the outcome.

**(b)** Example reward schedule. Reward contingency reversed every 8–12 trials, following a uniform distribution. The reward schedule was the same in the main experiment and the non-social control experiment.

**(c)** "Social influence" from computer agents on choice adjustment. Choice switch probability as a function of group consensus and direction (with vs. against) of the majority of the group. Results indicated a main effect of direction ($F_{1,41} = 95.63$, $p < 1.0 \times 10^{-11}$), a main effect of consensus ($F_{2,109} = 47.29$, $p < 1.0 \times 10^{-14}$), and an interaction effect ($F_{1,109} = 5.23$, $p = 0.024$). Solid lines indicate actual data (mean ± within-subject SEM). Shaded error bars represent the 95% highest density interval (HDI) of mean effects computed from the posterior predictive distribution of the winning model M1b (social model with only instantaneous social information but without social leaning).

**(d)** "Social influence" from computer agents on bet adjustments. Bet difference as a function of group consensus and direction (with vs. against) of the majority of the group. Results indicated a main effect of direction ($F_{1,140} = 40.44$, $p < 1.0 \times 10^{-8}$), a main effect of consensus ($F_{2,140} = 3.19$, $p = 0.044$), and no interaction effect ($F_{1,140} = 0.116$, $p = 0.734$). Format is as in Supplementary Fig. 1c.

**(e–f)** Non-improvement in performance after adjustment. (e) Accuracy of Choice 2 did not differ from that of Choice 1 ($t_{35} = 0.8798$, $p = 0.385$). (f) Magnitude of Bet 2 did not differ from that of Bet 1 ($t_{35} = 1.5421$, $p = 0.132$).

**(g)** Comparison of learning-related parameters between the main study and the non-social control study, both fitted with M6b, although M6b was not the winning model for the non-social control data set. Crucially, 95% HDI of the parameter related to $V_{other}$ in the non-social control study (blue bar) contained zero, and it was smaller than the same parameter of the main study (95% HDI of the difference: [−1.027, −0.123]). When all models were fitted to data from the non-social control study, M1b was the winning model, showing no evidence of social learning.

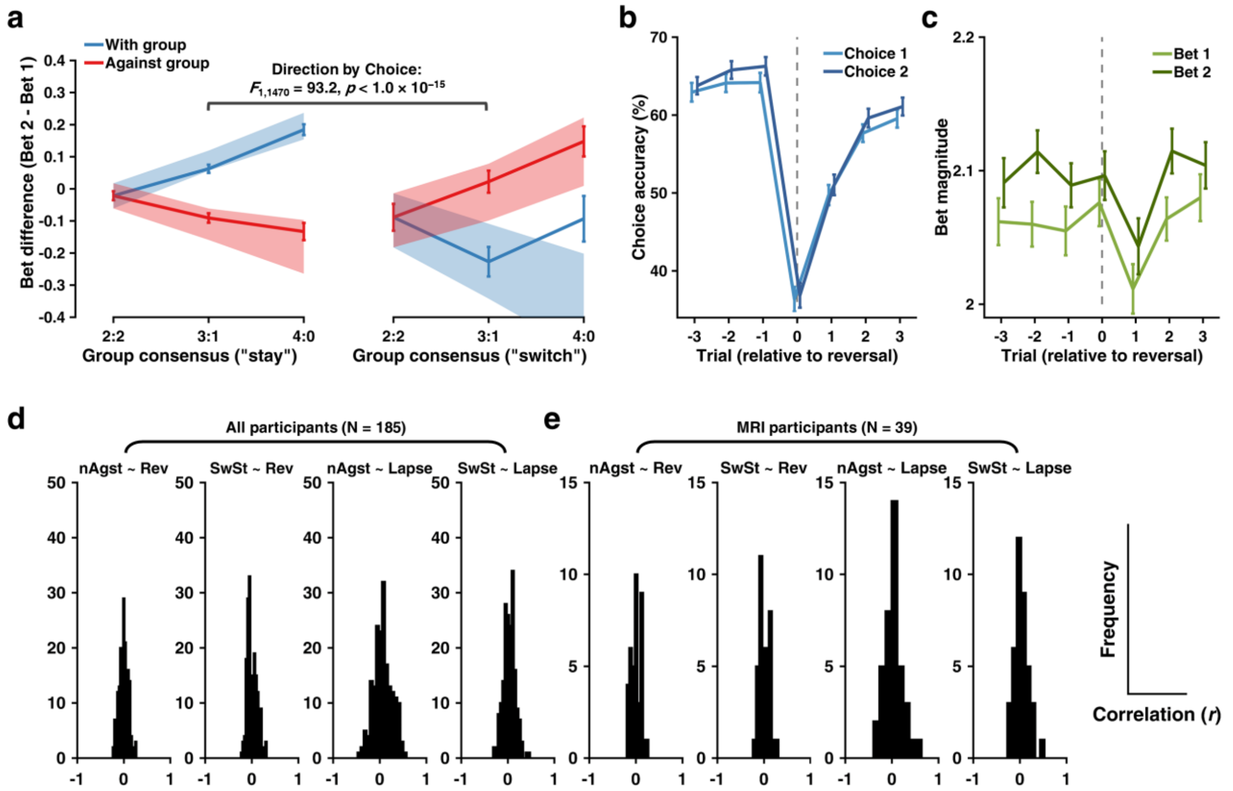


**Supplementary Figure 2**

**Behavioral data and their relationship to task structure.**

**(a)** Bet difference analyses separated by stay vs. switch trials. Bet difference analyses were performed with data separated by stay trials (C1 = C2, left panel) and switch trials (C1 ≠ C2, right panel). Results indicated a main effect of choice (stay vs. switch; $F_{1,1471} = 16.5$, $p < 1.0 \times 10^{-4}$), a two-way interaction between direction and choice ($F_{1,1470} = 93.2$, $p < 1.0 \times 10^{-15}$), and a two-way interaction between group consensus and choice ($F_{1,1456} = 12.5$, $p < 1.0 \times 10^{-5}$). All other effects were not significant (*p*s > 0.05). Solid lines indicate actual data (mean ± within-subject SEM). Shaded error bars represent the 95% highest density interval (HDI) of the mean effect computed from the winning model M6b's posterior density (posterior predictive checks). **(b)** Choice accuracy across reversal. Participants' choice accuracy as a function of choice type (Choice 1, light blue; Choice 2, dark blue) and trial positions relative to the reversal (−3, −2, −1, 0, 1, 2, 3). Results indicated a main effect of choice type ($F_{1,2392} = 4.45$, $p = 0.035$) and a main effect of position ($F_{6,2392} = 136.24$, $p < 1.0 \times 10^{-15}$; post-hoc pairwise comparison with Tukey's correction: "−3 vs. 0", $F_{1,2392} = 21.30$, $p < 1.0 \times 10^{-4}$; "−2 vs. 0", $F_{1,2392} = 22.57$, $p < 1.0 \times 10^{-4}$; "−1 vs. 0", $F_{1,2392} = 22.79$, $p < 1.0 \times 10^{-4}$; "+1 vs. 0", $F_{1,2392} = 10.97$, $p < 1.0 \times 10^{-4}$; "+2 vs. 0", $F_{1,2392} = 17.55$, $p < 1.0 \times 10^{-4}$; "+3 vs. 0", $F_{1,2392} = 18.88$, $p < 1.0 \times 10^{-4}$). No interaction was observed. All error bars indicate within-subject SEM. Dashed vertical line indicates the reversal.

**(c)** Bet magnitude across reversal. Participants' bet magnitude as a function of bet type (Bet 1, light green; Bet 2, dark green) and trials positions relative to the reversal (−3, −2, −1, 0, 1, 2, 3). Results indicated a main effect of bet type ($F_{1,184} = 7.10$, $p = 0.0083$) and a main effect of position ($F_{6,2208} = 3.52$, $p = 0.0018$; post-hoc pairwise comparison with Tukey's correction: "−2 vs. +1", $F_{1,2208} = 3.52$, $p = 0.008$; "+1 vs. 0", $F_{1,2208} = -3.47$, $p = 0.0097$; "+1 vs. +2", $F_{1,2208} = -3.65$, $p < 0.005$; "+1 vs. +3", $F_{1,2208} = -3.80$, $p = 0.0028$). Format is as in Supplementary Fig. 2b.

**(d–e)** Histograms of correlation coefficients between behaviors and task structure. Correlations were examined between preference-weighted dissenting social information and time of reversal (nAgst ~ Rev), preference-weighted dissenting social information and lapse error (nAgst ~ Lapse), switch-stay behavior and time of reversal (SwSt ~ Rev), switch-stay behavior and lapse error (SwSt ~ Lapse), both for the entire sample (d) and the MRI sample (e). Results indicated no pattern between these behaviors and task structure, as all histograms of correlation coefficients were stably centered around zero (two-tailed one sample *t*-tests, all *p*s > 0.05). Note that, the lapse error was defined as choosing one choice option on Choice 1 when the model strongly favored the alternative (modeled action probability >= 95%). For example, when the model predicted *p*(A) of Choice 1 was 95% (or higher) yet the participants actually chose option B, this trial was referred to as a lapse error. Inset: axis labels.

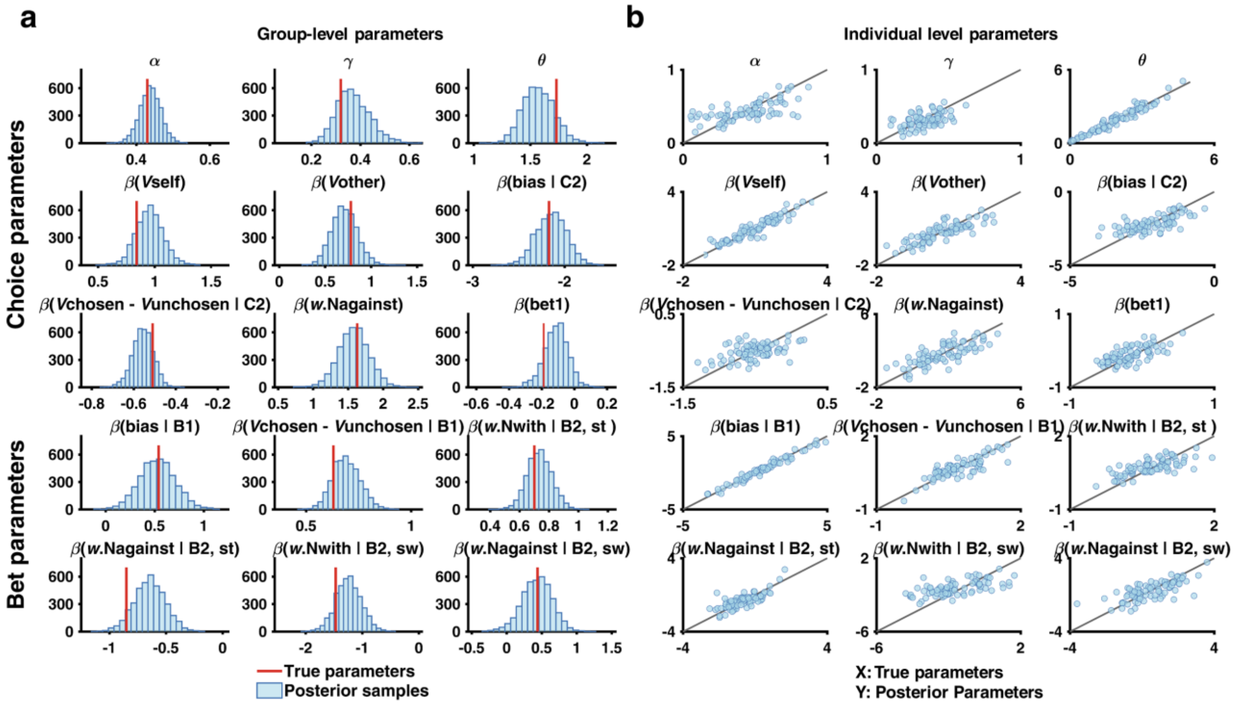


**Supplementary Figure 3**

**Parameter recovery.**

A random sample of group-level parameters was drawn from the winning model M6b's group-level joint posterior distribution (Fig 2f–i) to simulate 80 synthetic participants' behavioral data (Choice 1, Choice 2, Bet 1, Bet 2) for the social influence task. The data generating model M6b was fitted to data from these 80 synthetic participants, and parameter estimates were then compared to the data generating parameters, both at the group-level and at the individual level. All parameters could be accurately and selectively recovered, showing proper identifiability of model parameters: (a) for the group-level, "true" parameters falling within 95% HDI of each parameter's posterior density; (b) for the individual level, "true" parameters and estimated parameters were correlated along the identity line.

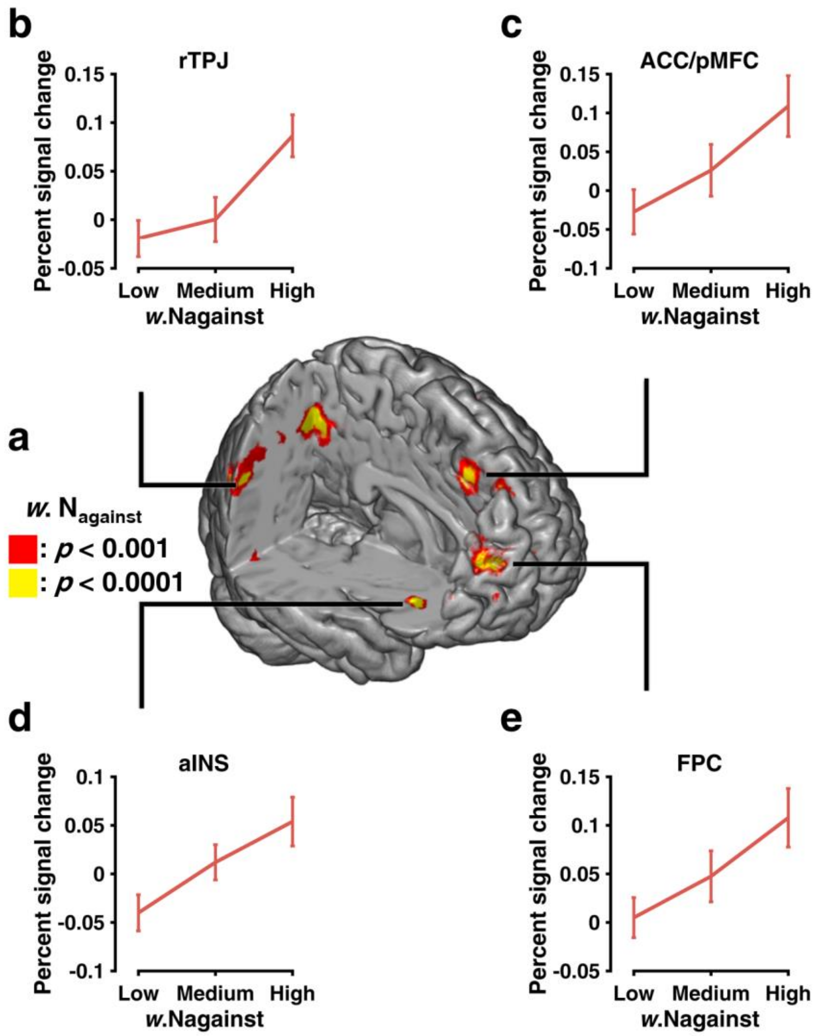


**Supplementary Figure 4**

**Neural substrates of dissenting social information.**

**(a)** Activities (red/yellow) in right temporal-parietal junction (rTPJ), anterior cingulate cortex (ACC)/posterior Medial Frontal Cortex (pMFC), bilateral anterior insula (aINS) and frontopolar cortex (FPC) scaled parametrically with the dissenting social information ($w.N_{against}$). Color coding indicates SPM thresholds corresponding to $p < 0.001$ uncorrected and $p < 0.0001$ uncorrected. Actual results were TFCE whole-brain FWE corrected at $p < 0.05$.

**(b–e)** Percent signal change (PSC) of BOLD time series in the corresponding ROIs for trials in which $w.N_{against}$ was low, medium, and high ($33^{rd}$, $66^{th}$, and $100^{th}$ percentile). All show a significant linear increasing trend (*F*-tests, *p*s < 0.001). For sub-figure d, we chose the right aINS because its supra-threshold cluster was larger than the left one (Supplementary Table 4). Using the left aINS yielded similar results. All error bars indicate SEM.

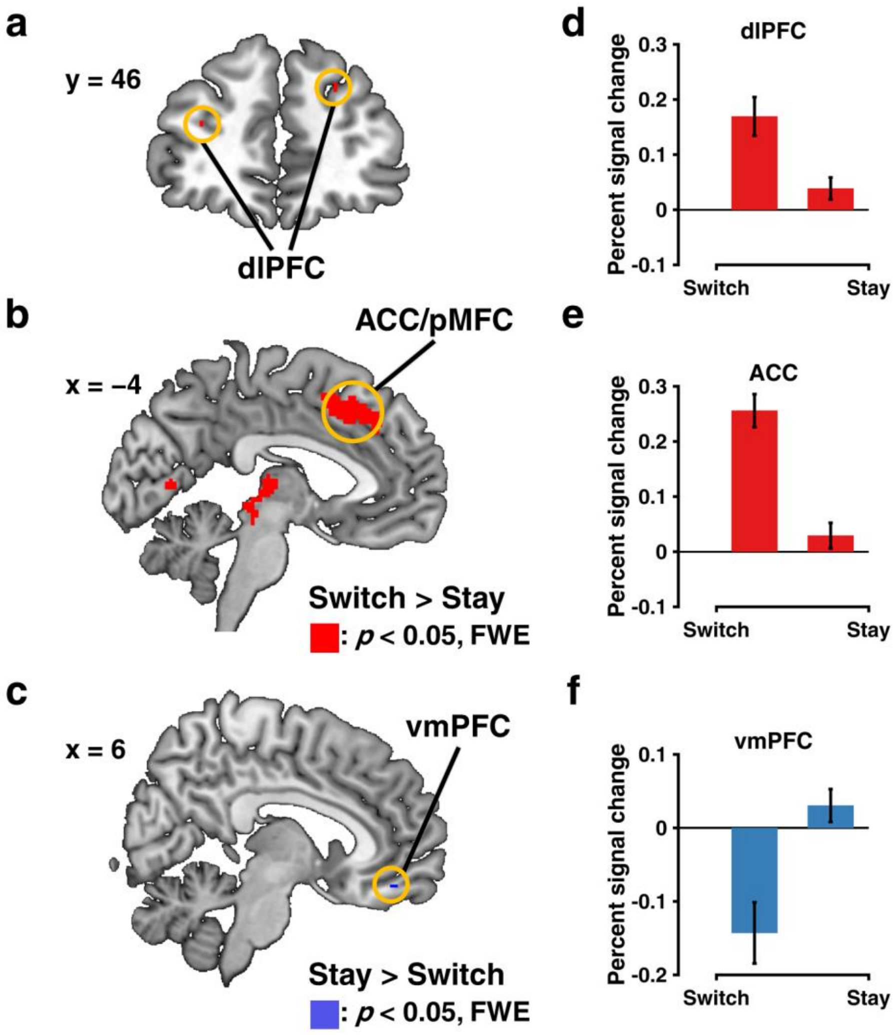


**Supplementary Figure 5**

**Neural substrates of choice adjustment.**

**(a–b)** Bilateral dorsolateral prefrontal cortex (dlPFC) and the anterior cingulate cortex (ACC)/posterior medial frontal cortex (pMFC) were more activated when individuals made a switch during choice adjustment. Color coding indicates SPM threshold corresponding to $p < 0.05$ FWE corrected. Actual results were TFCE whole-brain FWE corrected at $p < 0.05$.
**(c)** Ventromedial prefrontal cortex (vmPFC) was more activated when individuals made a stay during choice adjustment. Format is as in Supplementary Fig. 5a.
**(d–f)** Percent signal change (PSC) of BOLD time series in the corresponding ROIs for switching trials as opposed to staying trials. All show a significant difference (paired *t*-tests, *p*s < 0.001). For For sub-figure b, we chose the left dlPFC because its supra-threshold cluster was larger than the right one (Supplementary Table 4). Using the right dlPFC yielded similar results. All error bars indicate SEM.

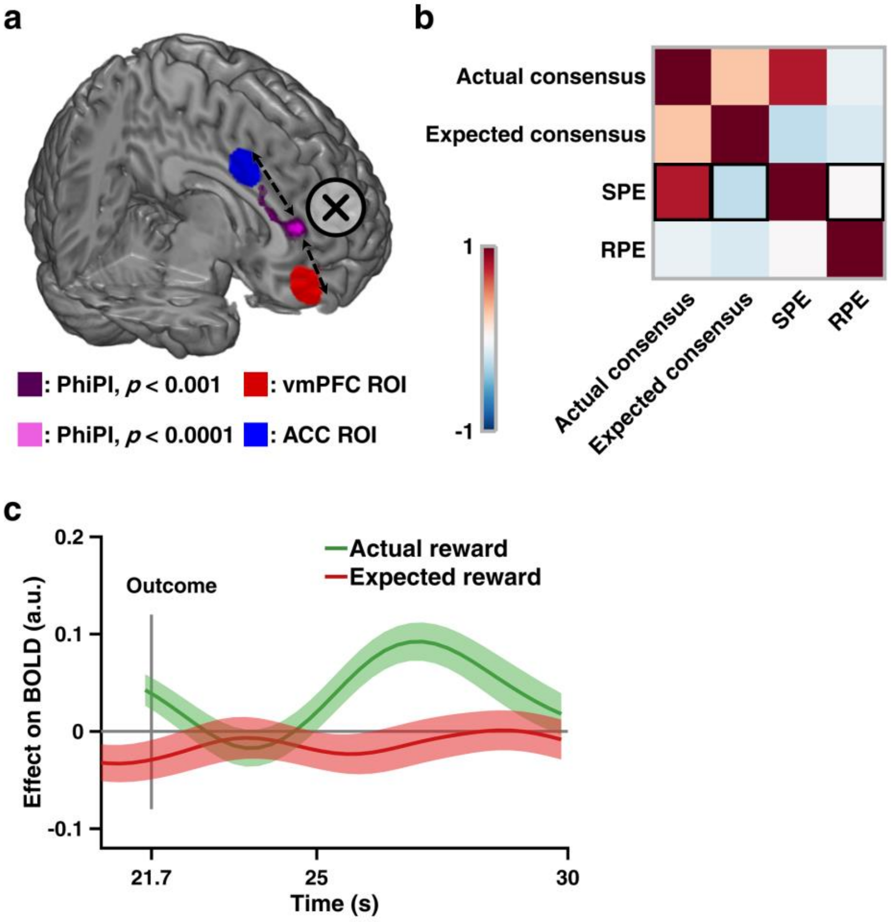


**Supplementary Figure 6**

**Functional connectivity.**

**(a)** Physio-physiological interaction (PhiPI) between ventromedial prefrontal cortex (vmPFC) and anterior cingulate cortex (ACC). Two seed regions, vmPFC (red) and ACC (blue) elicited connectivity activations (target regions) in the (rostral) medial prefrontal cortex (mPFC; magenta). Color coding indicates SPM thresholds corresponding to $p < 0.001$ uncorrected and $p < 0.0001$ uncorrected, respectively. Actual results were TFCE whole-brain FWE corrected at $p < 0.05$.

(**b**) Correlation matrix of variables related to social prediction error (SPE). Crucially, SPE and RPE were not correlated ($r = 0.0039$, $p = 0.431$).

**(c)** (Non-)Association between left putamen and components of reward prediction error (RPE). BOLD time series of the left putamen was only positively correlated with the actual reward outcome ($\beta = 0.0633$, $p < 1.0 \times 10^{-4}$, permutation test; green line), but not negatively correlated with the expected outcome ($\beta = -0.0137$, $p = 0.4854$, permutation test, red line). Lines and shaded areas show mean ± SEM of $\beta$ weights across participants.

#### Supplementary Tables

##### Supplementary Table 1

**Linear mixed-effect models for the measurement of choice switch probability and bet difference.**

| Model | AIC | *p* |
|---|---|---|
| Choice Switch Probability | | |
| y ~ dir*con + (1|sub) | −506.75 | -- |
| y ~ dir*con + (1|sub) + (1|gender) + (1|group) | −502.84 | .956 |
| y ~ dir*con + (1|sub) + (1|gender) + (1|group) + (1|dir:sub) ← | −526.13 | < .001 |
| y ~ dir*con + (1|sub) + (1|gender) + (1|group) + (1|con:sub) | −500.84 | 1.000 |
| y ~ dir*con + (1|sub) + (1|gender) + (1|group) + (1|dir:sub) + (1|con:sub) | −524.13 | < .001 |
| Bet Difference | | |
| y ~ dir*con + (1|sub) ← | 101.21 | -- |
| y ~ dir*con + (1|sub) + (1|gender) + (1|group) | 105.21 | 1.000 |
| y ~ dir*con + (1|sub) + (1|gender) + (1|group) + (1|dir:sub) | 107.21 | .983 |
| y ~ dir*con + (1|sub) + (1|gender) + (1|group) + (1|con:sub) | 107.21 | 1.000 |
| y ~ dir*con + (1|sub) + (1|gender) + (1|group) + (1|dir:sub) + (1|con:sub) | 109.21 | .937 |
| Bet Difference split by "stay" vs. "switch" trials | | |
| y ~ dir*con*StSw + (1|sub) ← | 2115 | -- |
| y ~ dir*con*StSw + (1|sub) + (1|gender) + (1|group) | 2120 | 1.000 |
| y ~ dir*con*StSw + (1|sub) + (1|gender) + (1|group) + (1|dir:sub) | 2121 | 1.000 |
| y ~ dir*con*StSw + (1|sub) + (1|gender) + (1|group) + (1|con:sub) | 2121 | 1.000 |
| y ~ dir*con*StSw + (1|sub) + (1|gender) + (1|group) + (1|dir:sub) + (1|con:sub) | 2123 | 1.000 |

Note: Models are specified in the R package "lme4" syntax: "A*B" denotes the main effect plus the interaction effect between variables A and B; "(1|var)" denotes the singular random effect; "(1|A:B)" denotes the interaction random effect between variables A and B. "dir" = direction (with vs. against the group). "con" = group consensus level (2:2, 3:1, 4:0). "sub" = subject. "gender" = participants' gender (female vs. male). "group" = measurement group (MRI vs. behavioral). "StSw" = trial types (stay vs. switch). AIC = Akaike Information Criterion; lower AIC value indicates higher model evidence. *p* values are calculated with $\chi^2$ tests. Models that best capture the variance structure of the data are indicated by a left-pointing arrow sign (←).

**Supplementary Table 2**

**Linear mixed-effect models for the measurement of choice accuracy and bet magnitude.**

| Model | AIC | *p* |
|---|---|---|
| Choice Accuracy | | |
| `y ~ type*pos + (1|sub)` ← | −1559.9 | -- |
| `y ~ type*pos + (1|sub) + (1|gender) + (1|group)` | −1555.9 | 1.000 |
| `y ~ type*pos + (1|sub) + (1|gender) + (1|group) + (1|type:sub)` | −1553.9 | 1.000 |
| Bet Magnitude | | |
| `y ~ type*pos + (1|sub)` | 713.16 | -- |
| `y ~ type*pos + (1|sub) + (1|gender) + (1|group)` | 712.83 | .115 |
| `y ~ type*pos + (1|sub) + (1|gender) + (1|group) + (1|type:sub)` ← | 656.75 | < .001 |

Note: Models are specified in the R package "lme4" syntax: "A*B" denotes the main effect plus the interaction effect between variables A and B; "(1|var)" denotes the singular random effect; "(1|A:B)" denotes the interaction random effect between variables A and B. "type" = choice type (1$^{st}$ choice vs. 2$^{nd}$ choice; or 1$^{st}$ bet vs. 2$^{nd}$ bet). "pos" = trial position relative to reversal (−3, −2, −1, 0, +1, +2, +3). "sub" = subject. "gender" = participants' gender (female vs. male). "group" = measurement group (MRI vs. behavioral). AIC = Akaike Information Criterion; lower AIC value indicates higher model evidence. *p* values are calculated with $\chi^2$ tests. Models that best capture the variance structure of the data are indicated by a left-pointing arrow sign (←).

**Supplementary Table 3**

**Specification of computational models.**

| Class | Model | Specification |
|---|---|---|
| | Commons | $\mathbb{V}_t = [V_t(\mathrm{A}), V_t(\mathrm{B})]$<br>$C1_t \sim Categorical(Softmax(\beta_{\mathrm{v}}\,\mathbb{V}_t))$<br>$U_{\mathrm{bet1},t} = \beta_{\mathrm{bias}_{B1}} + \beta_{\mathrm{vdiff}_{B1}}(V_{\mathrm{chosen},C1,t} - V_{\mathrm{unchosen},C1,t})$<br>$B1_t \sim OrderedLogistic(U_{\mathrm{bet1},t} \mid \theta)$<br>$V_t(\mathrm{switch}) = \beta_{\mathrm{bias}_{C2}} + \beta_{\mathrm{vdiff}_{C2}}(V_{\mathrm{chosen},C1,t} - V_{\mathrm{unchosen},C1,t})$<br>$C2 \sim Bernoulli(V_t(\mathrm{switch}))$<br>$U_{\mathrm{bet2},t} = \begin{cases} U_{\mathrm{bet1},t} + \beta_{\mathrm{bias}_{\mathrm{stay}}}, & \text{if } C1 = C2 \\ U_{\mathrm{bet1},t} + \beta_{\mathrm{bias}_{\mathrm{switch}}}, & \text{if } C1 \neq C2 \end{cases}$<br>$B2_t \sim OrderedLogistic(U_{\mathrm{bet2},t} \mid \theta)$ |
| Non-social models | M1a | $\delta_{\mathrm{chosen},C2,t} = R_t - V_{\mathrm{chosen},C2,t}$<br>$V_{\mathrm{chosen},C2,t+1} = V_{\mathrm{chosen},C2,t} + \alpha\delta_{\mathrm{chosen},C2,t}$<br>$V_{\mathrm{unchosen},C2,t+1} = V_{\mathrm{unchosen},C2,t}$ |
| | M1b | $\delta_{\mathrm{chosen},C2,t} = R_t - V_{\mathrm{chosen},C2,t}$<br>$\delta_{\mathrm{unchosen},C2,t} = -R_t - V_{\mathrm{unchosen},C2,t}$<br>$V_{\mathrm{chosen},C2,t+1} = V_{\mathrm{chosen},C2,t} + \alpha\delta_{\mathrm{chosen},C2,t}$<br>$V_{\mathrm{unchosen},C2,t+1} = V_{\mathrm{unchosen},C2,t} + \alpha\delta_{\mathrm{unchosen},C2,t}$ |
| | M1c | $\delta_{\mathrm{chosen},C2,t} = R_t - V_{\mathrm{chosen},C2,t}$<br>$V_{\mathrm{chosen},t+1} = V_{\mathrm{chosen},t} + k\alpha_t\delta_{\mathrm{chosen},C2,t}$<br>$V_{\mathrm{unchosen},C2,t+1} = V_{\mathrm{unchosen},C2,t}$<br>$\alpha_{t+1} = \lambda\lvert\delta_{\mathrm{chosen},t}\rvert + (1-\lambda)\,\alpha_t$ |

| | | |
|---|---|---|
| Social models: instantaneous social influence | Commons | $\begin{aligned} \mathbb{V}_t &= [V_t(\mathrm{A}), V_t(\mathrm{B})] \\ C1_t &\sim \mathit{Categorical}(\mathit{Softmax}(\beta_{\mathrm{v}}\, \mathbb{V}_t)) \\ U_{\mathrm{bet1},t} &= \beta_{\mathrm{bias}_{B1}} + \beta_{\mathrm{vdiff}_{B1}}(V_{\mathrm{chosen},C1,t} - V_{\mathrm{unchosen},C1,t}) \\ B1_t &\sim \mathit{OrderedLogistic}(U_{\mathrm{bet1},t} \mid \theta) \\ w.\mathrm{N}_{\mathrm{against},t} &= \frac{\sum_{s=1}^{K} w_{s,t}}{\sum_{s=1}^{4} w_{s,t}}, K = 0,1,...,4 \\ w.\mathrm{N}_{\mathrm{with},t} &= \frac{\sum_{s=1}^{4-K} w_{s,t}}{\sum_{s=1}^{4} w_{s,t}} \\ V_t(\mathrm{switch}) &= \beta_{\mathrm{bias}_{C2}} + \beta_{\mathrm{vdiff}_{C2}}(V_{\mathrm{chosen},C1,t} - V_{\mathrm{unchosen},C1,t}) + \beta_{\mathrm{against}}\, w.\mathrm{N}_{\mathrm{against},t} \\ C2 &\sim \mathit{Bernoulli}(V_t(\mathrm{switch})) \\ U_{\mathrm{bet2},t} &= \begin{cases} U_{\mathrm{bet1},t} + \beta_{\mathrm{with}_{\mathrm{stay}}}\, w.\mathrm{N}_{\mathrm{with},t} + \beta_{\mathrm{against}_{\mathrm{stay}}}\, w.\mathrm{N}_{\mathrm{against},t} & \text{, if } C1 = C2 \\ U_{\mathrm{bet1},t} + \beta_{\mathrm{with}_{\mathrm{switch}}}\, w.\mathrm{N}_{\mathrm{with},t} + \beta_{\mathrm{against}_{\mathrm{switch}}}\, w.\mathrm{N}_{\mathrm{against},t}, & \text{if } C1 \neq C2 \end{cases} \\ B2_t &\sim \mathit{OrderedLogistic}(U_{\mathrm{bet2},t} \mid \theta) \end{aligned}$ |
| | M2a | $\begin{aligned} \delta_{\mathrm{chosen},C2,t} &= R_t - V_{\mathrm{chosen},C2,t} \\ V_{\mathrm{chosen},C2,t+1} &= V_{\mathrm{chosen},C2,t} + \alpha\delta_{\mathrm{chosen},C2,t} \\ V_{\mathrm{unchosen},C2,t+1} &= V_{\mathrm{unchosen},C2,t} \end{aligned}$ |
| | M2b | $\begin{aligned} \delta_{\mathrm{chosen},C2,t} &= R_t - V_{\mathrm{chosen},C2,t} \\ \delta_{\mathrm{unchosen},C2,t} &= -R_t - V_{\mathrm{unchosen},C2,t} \\ V_{\mathrm{chosen},C2,t+1} &= V_{\mathrm{chosen},C2,t} + \alpha\delta_{\mathrm{chosen},C2,t} \\ V_{\mathrm{unchosen},C2,t+1} &= V_{\mathrm{unchosen},C2,t} + \alpha\delta_{\mathrm{unchosen},C2,t} \end{aligned}$ |
| | M2c | $\begin{aligned} \delta_{\mathrm{chosen},C2,t} &= R_t - V_{\mathrm{chosen},C2,t} \\ V_{\mathrm{chosen},t+1} &= V_{\mathrm{chosen},t} + k\alpha_t\delta_{\mathrm{chosen},C2,t} \\ V_{\mathrm{unchosen},C2,t+1} &= V_{\mathrm{unchosen},C2,t} \\ \alpha_{t+1} &= \lambda\lvert\delta_{\mathrm{chosen},t}\rvert + (1-\lambda)\alpha_t \end{aligned}$ |

| | | |
|---|---|---|
| Social models: instantaneous social influence & social learning | Commons | $\mathbb{V}_{\text{self},t} = [V_{\text{self},t}(\text{A}), V_{\text{self},t}(\text{B})]$<br>$\mathbb{V}_{\text{other},t} = [V_{\text{other},t}(\text{A}), V_{\text{other},t}(\text{B})]$<br>$\mathbb{V}_{t} = \beta_{\text{vself}}\,\mathbb{V}_{\text{self},t} + \beta_{\text{vother}}\,\mathbb{V}_{\text{other},t}$<br>$C1_{t} \sim Categorical(Softmax(\mathbb{V}_{t}))$<br>$U_{\text{bet1},t} = \beta_{\text{bias}B1} + \beta_{\text{vdiff}B1}(V_{\text{chosen},C1,t} - V_{\text{unchosen},C1,t})$<br>$B1_{t} \sim OrderedLogistic(U_{\text{bet1},t} \mid \theta)$<br>$w.\text{N}_{\text{against},t} = \frac{\sum_{s=1}^{K} w_{s,t}}{\sum_{s=1}^{4} w_{s,t}}, K = 0,1,...,4$<br>$w.\text{N}_{\text{with},t} = \frac{\sum_{s=1}^{4-K} w_{s,t}}{\sum_{s=1}^{4} w_{s,t}}$<br>$V_{t}(\text{switch}) = \beta_{\text{bias}C2} + \beta_{\text{vdiff}C2}(V_{\text{chosen},C1,t} - V_{\text{unchosen},C1,t}) + \beta_{\text{against}}\,w.\text{N}_{\text{against},t}$<br>$C2 \sim Bernoulli(V_{t}(\text{switch}))$<br>$U_{\text{bet2},t} = \begin{cases} U_{\text{bet1},t} + \beta_{\text{with}_{\text{stay}}}\,w.\text{N}_{\text{with},t} + \beta_{\text{against}_{\underline{\text{stay}}}}\,w.\text{N}_{\text{against},t} & \text{, if } C1 = C2 \\ U_{\text{bet1},t} + \beta_{\text{with}_{\text{switch}}}\,w.\text{N}_{\text{with},t} + \beta_{\text{against}_{\text{switch}}}\,w.\text{N}_{\text{against},t}, & \text{if } C1 \neq C2 \end{cases}$<br>$B2_{t} \sim OrderedLogistic(U_{\text{bet2},t} \mid \theta)$<br>$\Phi(x) = \frac{1}{1+e^{-x}}$<br>$\delta_{\text{self},\text{chosen},C2,t} = R_{\text{self},t} - V_{\text{self},\text{chosen},C2,t}$<br>$\delta_{\text{self},\text{unchosen},C2,t} = -R_{\text{self},t} - V_{\text{self},\text{unchosen},C2,t}$<br>$V_{\text{self},\text{chosen},C2,t+1} = V_{\text{self},\text{chosen},C2,t} + \alpha\delta_{\text{self},\text{chosen},C2,t}$<br>$V_{\text{self},\text{unchosen},C2,t+1} = V_{\text{self},\text{unchosen},C2,t} + \alpha\delta_{\text{self},\text{unchosen},C2,t}$ |
| | M3 | $\delta_{s,\text{chosen},C2,t} = R_{s,t} - V_{s,\text{chosen},C2,t}, s = 1,2,3,4$<br>$\delta_{s,\text{unchosen},C2,t} = -R_{s,t} - V_{s,\text{unchosen},C2,t}$<br>$V_{s,\text{chosen},C2,t+1} = V_{s,\text{chosen},C2,t} + \alpha_{o}\delta_{s,\text{chosen},C2,t}$<br>$V_{s,\text{unchosen},C2,t+1} = V_{s,\text{unchosen},C2,t} + \alpha_{o}\delta_{s,\text{unchosen},C2,t}$<br>$V_{\text{other},t+1}(\text{A}) = \sum_{s=1}^{4} w_{s,t} V_{s,t+1}(\text{A})$<br>$V_{\text{other},t+1}(\text{B}) = \sum_{s=1}^{4} w_{s,t} V_{s,t+1}(\text{B})$<br>$V_{\text{other},t+1}(\text{A}) = 2\Phi(V_{\text{other},t+1}(\text{A})) - 1$<br>$V_{\text{other},t+1}(\text{B}) = 2\Phi(V_{\text{other},t+1}(\text{B})) - 1$ |

| | |
|---|---|
| M4 | $\rho_{s,t}(\mathrm{A}) = I_{0.5}(1+\sum_{t=T-2}^{T} C2_{\mathrm{B},s,t}, 1+\sum_{t=T-2}^{T} C2_{\mathrm{A},s,t})$<br>$\rho_{s,t}(\mathrm{B}) = 1-\rho_{s,t}(\mathrm{A})$<br>$V_{\mathrm{other},t+1}(\mathrm{A}) = \sum_{s=1}^{4} w_{s,t}\rho_{s,t}(\mathrm{A})$<br>$V_{\mathrm{other},t+1}(\mathrm{B}) = \sum_{s=1}^{4} w_{s,t}\rho_{s,t}(\mathrm{B})$<br>$V_{\mathrm{other},t+1}(\mathrm{A}) = 2\Phi(V_{\mathrm{other},t+1}(\mathrm{A}))-1$<br>$V_{\mathrm{other},t+1}(\mathrm{B}) = 2\Phi(V_{\mathrm{other},t+1}(\mathrm{B}))-1$ |
| M5 | $V_{\mathrm{other},t+1}(\mathrm{A}) = \sum_{s=1}^{K_{\mathrm{A}}} w_{s,t}R_{s,t}, K_{\mathrm{A}} = 0,1,...,4$<br>$V_{\mathrm{other},t+1}(\mathrm{B}) = \sum_{s=1}^{4-K_{\mathrm{A}}} w_{s,t}R_{s,t}$<br>$V_{\mathrm{other},t+1}(\mathrm{A}) = 2\Phi(V_{\mathrm{other},t+1}(\mathrm{A}))-1$<br>$V_{\mathrm{other},t+1}(\mathrm{B}) = 2\Phi(V_{\mathrm{other},t+1}(\mathrm{B}))-1$ |
| M6a | $V_{\mathrm{other},t+1}(\mathrm{A}) = \sum_{s=1}^{K_{\mathrm{A}}}\sum_{t=T-2}^{T} w_{s,t}\gamma^{T-i}R_{s,i}, K_{\mathrm{A}} = 0,1,...,4$<br>$V_{\mathrm{other},t+1}(B) = \sum_{s=1}^{4-K_{\mathrm{A}}}\sum_{t=T-2}^{T} w_{s,t}\gamma^{T-i}R_{s,i}$<br>$V_{\mathrm{other},t+1}(\mathrm{A}) = 2\Phi(V_{\mathrm{other},t+1}(\mathrm{A}))-1$<br>$V_{\mathrm{other},t+1}(\mathrm{B}) = 2\Phi(V_{\mathrm{other},t+1}(\mathrm{B}))-1$ |
| M6b | $V_t(\mathrm{switch}) = \beta_{\mathrm{bias}C2} + \beta_{\mathrm{vdiff}C2}(V_{\mathrm{chosen},C1,t} - V_{\mathrm{unchosen},C1,t}) + \beta_{\mathrm{against}} w.\mathrm{N}_{\mathrm{against},t} + \beta_{\mathrm{b1}}\mathrm{Bet1}_t$<br>$V_{\mathrm{other},t+1}(\mathrm{A}) = \sum_{s=1}^{K_{\mathrm{A}}}\sum_{t=T-2}^{T} w_{s,t}\gamma^{T-i}R_{s,i}, K_{\mathrm{A}} = 0,1,...,4$<br>$V_{\mathrm{other},t+1}(B) = \sum_{s=1}^{4-K_{\mathrm{A}}}\sum_{t=T-2}^{T} w_{s,t}\gamma^{T-i}R_{s,i}$<br>$V_{\mathrm{other},t+1}(\mathrm{A}) = 2\Phi(V_{\mathrm{other},t+1}(\mathrm{A}))-1$<br>$V_{\mathrm{other},t+1}(\mathrm{B}) = 2\Phi(V_{\mathrm{other},t+1}(\mathrm{B}))-1$ |

Note: A/B = choice option. C1 = Choice 1; B1 = Bet 1; C2 = Choice 2; B2 = Bet 2; $V$ = action value; $t$ = trial index; $\beta$s = beta weights; $U$ = bet utility; $\theta$ = threshold in ordered logistic models; $R$ = reward; $\alpha$ = learning rate in RL models; $\delta$ = reward prediction error in RL models; $k$ = weight of associability in the Pearce-Hall model; $\lambda$ = weight between prediction error and associability in the Pearce-Hall model; $w$ = preference weighting; $K$ = amount of opposite C1s from others; $\alpha_o$ = other-oriented learning rate in RL models; s = index of others; $\rho$ = option preference based on past actions; $I_{0.5}$(a, b) = cumulative distribution function of the beta distribution at the value of 0.5; $K_{\mathrm{A}}$ = amount of others choosing A for their C2; $\gamma$ = decay rate of others' reward history.

#### Supplementary Note 1

##### Procedure of the non-social control study

We carried out a non-social control study to determine whether participants' learning behavior in our main study was specific to a social situation. Participants in the control study underwent the decision-making task with four intelligent computer agents. We intentionally did so, instead of using four randomly behaving computer agents, in order to clarify if the social learning component in our main study was specific when participants were playing with human partners (Boorman et al., 2013). The procedure of the control study was largely similar to the one of the main study, except that there was no preference giving, because this would otherwise make participants feel unrealistic to rate computer agents. The control study was composed of a training phase and a testing phase, and they were on a single test day.

*Training phase*

On the test day, participants came to the behavioral lab to take part in the experiment. Upon arrival, they received the written task instruction and the consent form. After returned the written consent, participants were taken through a step-by-step task instruction by the experimenter and were given opportunities to ask any questions related to the task. Crucially, participants were instructed that they were about to play with four intelligent computer agents that could use an intelligent algorithm to learn the outcome contingency through trial-and-error, as well as consider other computer agents and the participants' choice to improve their performance. Unknown to the participants, the choice sequences were in fact obtained from a subset of participants (with similar task performance) from our main study, which entailed human learning. Additionally, participants were explicitly informed (a) what probabilistic reward meant by receiving examples, (b) that there was neither cooperation nor competition with the computer agents in this experiment, and (c) that the reward probability could reverse multiple times over the course of the experiment, but participants were not informed about when and how often this reversal would take place. To further enhance participants' motivation, we informed them that the amount they would gain from the experiment would be added to their base payment (see below).

The training phase consisted of 10 trials and differed from the testing phase in two aspects. Firstly, it used a different set of stimuli than those used in the testing phase to avoid any learning effect. Secondly, participants were given a longer response window to fully understand every step of the task. Specifically, each trial began with the stimuli presentation of two choice alternatives and participants were asked to decide on their Choice 1 (4000 ms) and Bet 1 (3000 ms). Then, all Choice 1 from four computer agents were displayed below the computer icons (3000 ms). Participants were then asked to adjust their choice (Choice 2; 4000 ms) and their bet (Bet 2; 3000 ms). Finally, outcomes of the participants were released (3000 ms), followed by a jittered inter-trial interval (ITI, 2000–4000 ms). To help participants familiarize themselves, we orally instructed them what to do on each step for the first two to three trials.

*Testing phase*

The testing phase took place immediately after the training phase. The testing phase consisted of 100 trials and used a different pair of stimuli from the training task (Supplementary Fig. 1a). Specifically, each trial began with the stimuli presentation of two choice alternatives and participants were asked to decide on their Choice 1 (3000 ms) and Bet 1 (2000 ms). Then, all Choice 1 from four computer agents were displayed below the computer icons (3000 ms). Participants were then asked to adjust their choice (Choice 2; 3000 ms) and their bet (Bet 2; 2000 ms). Finally, outcomes of the participants were released (3000 ms), followed by a jittered inter-trial interval (ITI, 2000–4000 ms). The entire procedure (training + testing) lasted about 2 hours.

*Reward payment*

Participants were compensated with a base payment of 20 Euro plus the reward they had achieved during the testing phase. In the testing phase, to prevent participants from careless responses on their Choice 1, they were explicitly instructed that on each trial, either their Choice 1 or their Choice 2 would be used to determine the final payoff. However, this did not affect the outcome delivery on the screen. Namely, although on some trials participants' Choice 1 was used to determine their payment, only outcomes that corresponded to their Choice 2 appeared on the screen. Additionally, when their total outcome was negative, no money was deducted from their final payment. Overall, participants gained 3.81 ± 3.76 Euro after completing the experiment. Finally, the experiment ended with an informal debriefing session.

**Supplementary Note 2**

**Parameter recovery**

Considering that there were multiple free parameters in the winning model, we verified whether parameters were identifiable using parameter recovery after the model fitting. Denoting $\theta$ as a generic parameter, we iterated the following:

(a) We randomly drew a set of group-level parameters from the joint posterior group-level distribution of M6b. That is, a group-level mean ($\mu_\theta$) and a group-level standard deviation ($\sigma_\theta$) of the parameter $\theta$. We repeated this procedure for all group-level parameters of the winning model M6b:

$$\mu_\theta, \sigma_\theta \sim p(\mu_\theta, \sigma_\theta \mid D). \tag{30}$$

(b) Next, we simulated 80 synthetic participants, whose parameters were randomly drawn from this set of group-level parameters. That is, individual-level parameters $\theta_i$ for 80 synthetic participants were sampled from the corresponding group-level parameters ($\mu_\theta, \sigma_\theta$) with a normal distribution. We repeated this procedure for all individual-level parameters of the winning model M6b:

$$\theta_i \sim Normal(\mu_\theta, \sigma_\theta). \tag{31}$$

(c) Then, we used the model (M6b) as a generative tool to simulate behavioral data for our social influence task, namely, to simulate Choice 1, Choice 2, Bet 1, and Bet 2 for 100 trials per participant. Individuals' choices and bets (denoted as $\hat{D}_i$) were sampled from the sampling distribution conditional on individual-level parameters ($\theta_i$) from the previous step (i.e., likelihood function):

$$\hat{D}_i \sim p(\hat{D} \mid \theta_i). \tag{32}$$

(d) We fit M6b to the simulated data ($\hat{D}_i$) in the same way as we did for the real data (D). That is, we fit M6b to the simulated individual data ($\hat{D}_i$) with MCMC using Stan, and obtained parameter estimates (i.e., posterior distributions) at both the group-level (e.g., $\mu_\theta^{'}$, $\sigma_\theta^{'}$) and individual level (e.g., $\theta_i^{'}$).

(e) Finally, we compared whether the posterior distributions at both the group-level and the individual level given the simulated data recovered the actual data generating parameters that were used to simulate those data, sampled from steps (a) and (b) (Supplementary Fig. 3).
